# A novel enhancer that regulates *Bdnf* expression in developing neurons

**DOI:** 10.1101/2021.11.18.469096

**Authors:** Emily Brookes, Ho Yu Alan Au, Wazeer Varsally, Christopher Barrington, Suzana Hadjur, Antonella Riccio

## Abstract

Brain derived neurotrophic factor (BDNF) is a critical secreted peptide that promotes neuronal differentiation and survival, and its downregulation is implicated in many neurological disorders. Here, we investigated the regulation of the mouse *Bdnf* gene in cortical neurons and identified a novel enhancer that promotes the expression of many *Bdnf* transcript variants during differentiation, increasing total *Bdnf* mRNA levels. Enhancer activity contributes to Bdnf-mediated effects on neuronal clustering and activity-dependent dendritogenesis. During *Bdnf* activation, enhancer-promoter contacts increase, and the region moves away from the repressive nuclear periphery. Our findings suggest that changes in nuclear structure may contribute to the expression of essential growth factors during neuronal development.

## Introduction

The *Brain Derived Neurotrophic Factor* (*BDNF*) gene encodes a neurotrophin with critical roles in brain development and functions, ranging from neuronal survival and differentiation during early development, to long-term potentiation and synaptic plasticity in the adult brain (Park and Poo, 2013). Aberrant *BDNF* expression has been implicated in a host of neurological diseases, including neuropsychiatric disorders such as schizophrenia (Di Carlo et al., 2019), stress (Notaras and van den Buuse, 2020) and depression (Caviedes et al., 2017); neurodegenerative diseases including Huntington’s (Yu et al., 2018; Zuccato and Cattaneo, 2007) and Alzheimer’s disease (Tanila, 2017); and neurodevelopmental disorders such as Rett syndrome (Li and Pozzo-Miller, 2014) and attention deficit hyperactivity disorder (ADHD) (Liu et al., 2015). Conversely, enhanced *BDNF* expression is linked to the neuroprotective effects of environmental enrichment (Dandi et al., 2018; Novkovic et al., 2015), exercise (Cotman et al., 2007; Wrann et al., 2013) and anti-depressants (Bjorkholm and Monteggia, 2016). Overexpression of *Bdnf* ameliorates symptoms in animal models of Rett syndrome (Chang et al., 2006; Sampathkumar et al., 2016) and Huntingdon’s disease (Canals et al., 2004; Gharami et al., 2008), and its neurotrophic effects suggest this may apply to other neurological disorders. Given the myriads of functions identified for *BDNF*, interrogating the regulation of the *BDNF* gene in neurons is of paramount importance to understand both developmental and disease mechanisms.

Rodent and human *BDNF* gene structure is complex, consisting of multiple 5’ exons, each containing its own promoter and 5’ untranslated region (5’UTR) that are alternatively spliced to a universal coding exon (Aid et al., 2007; Pruunsild et al., 2007; Timmusk et al., 1993) (**Fig. 1A**). Despite being translated into identical proteins, *Bdnf* mRNA variants exhibit specific expression patterns and physiological effects (Hallock et al., 2019; Hill et al., 2016; Maynard et al., 2016; Maynard et al., 2018; McAllan et al., 2018; Sakata and Duke, 2014; Sakata et al., 2013; Sakata et al., 2009). For example, disruption of exon I or II, but not IV or VI, enhances male aggression (Maynard et al., 2016) and impairs female maternal care (Maynard et al., 2018). Our current understanding of *Bdnf* gene control is centred on the distinct role of each promoter, however regulation through distal elements remains unclear.

**Figure 1.**
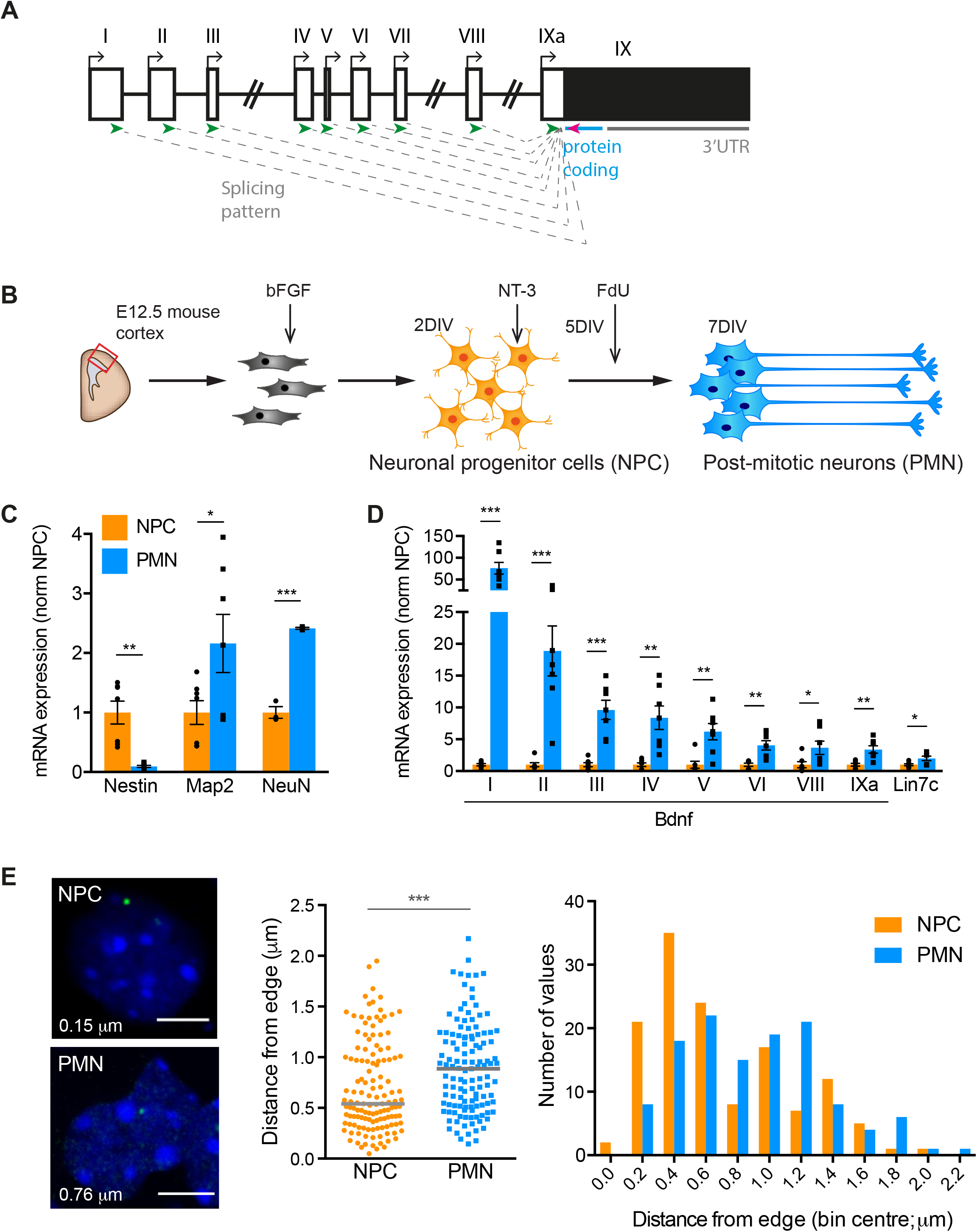
Expression of *Bdnf* isoforms increases over embryonic cortical development concomitantly with movement of the gene locus away from the nuclear periphery. **A)** Schematic of the *Bdnf* gene indicating the upstream exons (open boxes) which encode alternative 5’ untranslated regions (UTRs) that alternatively splice to the universal common exon (IX, black box) which encodes the protein coding sequence (blue) and the 3’ UTRs. Arrows indicate the position of the primers used to assess variant expression (green=forward, magenta=reverse). **B)** Schematic of the cell culture and differentiation paradigm of neuronal precursor cell (NPC) into post-mitotic neurons (PMN). bFGF, basic fibroblast growth factor. NT-3, neurotrophin-3. FdU, 5-fluoro-2’-deoxyuridine. DIV, days *in vitro*. **C)** Expression profile of an NPC-marker, *Nestin*, and neuronal markers, Map2 and NeuN, in NPCs and PMNs, assessed by qRT-PCR and normalised to NPC. Bars represent mean ± SEM, and points show results from different biological replicates (n = 7). ^*^*p* < 0.05, ^**^*p* < 0.01, ^***^*p* < 0.001; unpaired t-test (two-tailed). Nestin *p*=0.0005, *t*=4.707, *df*=12; Map2 p=0.0481, t=2.201, *df*=12, NeuN *p*=0.0001, t=14.15, *df*=4. **D)** Expression of *Bdnf* transcription variants and the downstream gene *Lin7c* during differentiation of NPC to PMN assessed by qRT-PCR and normalised to NPC. Bars represent mean ± SEM, and points show results from different biological replicates (n = 7). ^*^p < 0.05, ^**^p < 0.01, ^***^p < 0.001, ^****^p < 0.0001, unpaired t-test (two-tailed). Exon I *p*=0.0001, *t*=5.627, *df*=12; Exon II *p*=0.0007, *t*=4.549, *df*=12; Exon III *p*=0.0001, *t*=5.614, *df*=12; Exon IV *p*=0.0019, *t*=3.972, *df*=12; Exon V *p*=0.0029, *t*=3.734, *df*=12; Exon VI *p*=0.0021, *t*=3.899, *df*=12; Exon VIII *p*=0.0397, *t*=2.307, *df*=12; Exon IXa *p*=0.0023, *t*=3.846, *df*=12; *Lin7c p*=0.0248, *t*=2.565, *df*=12. **E)** Relocation of the *Bdnf* gene during neuronal development assessed by DNA-FISH combined with measurements of the distance of the signal from the edge of the nucleus. Left panel; representative images of confocal sections of DNA FISH showing nuclear localisation of *Bdnf* loci (green) in NPCs and PMNs. Nuclei were stained with DAPI (blue). For each image, the distance between the centre of the FISH signal and the edge of the nucleus is indicated. Scale bars, 2 μm. Middle panel; scatter dot plot of the distribution of the distance between *Bdnf* locus and the edge of the DAPI staining. Solid grey lines denote medians. ****p < 0.001 (p=0.0002), Mann-Whitney test (two-tailed). n = 133 (NPC), 123 (PMN) foci across 4 biological replicates. Right panel, histogram analysis of the same data measuring the distance between *Bdnf* locus and the edge of the DAPI staining in NPC (orange bars) and PMN (blue bars).

Enhancers are short regions of regulatory DNA, whose activity promotes the expression of their target gene(s) (Banerji et al., 1981). Combinations of enhancer elements confer spatially and temporally precise gene expression profiles (Carullo and Day, 2019). In linear chromosomal distance enhancers are often located far from the genes that they control, although within the three dimensional (3D) nuclear space they become proximal through enhancer-promoter looping (Schoenfelder and Fraser, 2019). Putative enhancers for *Bdnf* have been identified based on 3D proximity to the gene and H3K27ac occupancy (Beagan et al., 2020), and an intronic enhancer regulating both basal and stimulus-dependent expression of *Bdnf* was recently found for transcripts expressed from promoters I-III (Tuvikene et al., 2021).

Enhancer-promoter proximity can be critical for appropriate gene expression (Bartman et al., 2016; Deng et al., 2012; Greenwald et al., 2019; Kim et al., 2019; Morgan et al., 2017). Enhancer-promoter looping is supported by the genome architecture of the region and can occur in the context of topologically associated domains (TADs), which are megabase-sized regions of DNA that interact more commonly with each other than with the surrounding regions (Brookes and Riccio, 2019). Genome topology and gene activation is also affected by nuclear compartmentalisation, and therefore the position of the gene with respect to nuclear landmarks is important.

Here, we identify a novel enhancer region that loops to the *Bdnf* gene in neurons and is critical for *Bdnf* expression during neuronal differentiation and dendritic growth. We also show that the *Bdnf* gene is located in a previously undescribed sub-TAD and that the gene is repositioned away from the nuclear periphery during developmental upregulation. Together our results identify a mechanism of regulation that is implicated in *Bdnf* expression during both neurodevelopmental processes and neurological disease.

## Results

### Nuclear relocation of the activated *Bdnf* gene during neuronal differentiation

To study neuronal differentiation using a tractable model system (Nitarska et al., 2016), neurons were dissected from E12.5 mouse cortices and cultured with fibroblast growth factor (FGF) for 2 days *in vitro* (DIV) to generate a homogenous population of neuronal progenitor cells (NPCs) (**Fig. 1B**). After a few days, NPCs were differentiated into neurons by adding neurotrophin-3 (NT-3) and the anti-mitotic agent 5-fluoro-2’-deoxyuridine (Fdu) to remove remaining proliferating cells. Post-mitotic neurons (PMN) were harvested after 7 DIV. Expression analysis of the NPC-marker *Nestin* and the neuronal markers *Map2* and *NeuN* confirmed the reliability of this model system (**Fig. 1C**). The expression of different isoforms of *Bdnf* was analysed by quantitative reverse transcription PCR (qRT-PCR) with a reverse primer complementary to universal exon IX, and forward primers matching each 5’UTR (**Fig. 1A**). The expression of all *Bdnf* isoforms significantly increased during the differentiation of NPC to PMN, with the exon I-containing isoform showing the most substantial increase (**Fig. 1D**). The nearest downstream gene, *Lin7c*, also showed a significant increase from NPC to PMN (**Fig. 1D**).

To understand the mechanisms that facilitate the striking increase in *Bdnf* expression during neuronal differentiation, we first investigated the 3D nuclear position of the *Bdnf* gene. The nucleus is highly organised, with the nuclear periphery enriched in heterochromatin and providing a suitable environment to repress transcription (Brookes and Riccio, 2019). Movement away from the lamina is therefore often concomitant with either increased gene expression or increased competency for later expression (Peric-Hupkes et al., 2010). The *Bdnf* gene is known to relocate from the nuclear periphery to the nuclear interior during activation in response to kainate-induced seizures in the adult brain (Walczak et al., 2013). To understand whether this occurs during neuronal differentiation, DNA Fluorescence *In Situ* Hybridisation (DNA-FISH) was performed on both NPCs and PMNs using a BAC (Bacterial Artificial Chromosome) spanning the *Bdnf* locus. When the distance of *Bdnf* from the edge of the nucleus stained with 4⍰, 6-diamidino-2-phenylindole (DAPI) was measured, we observed significant movement of the locus away from the nuclear periphery (**Fig. 1E**).

### *Bdnf* loops to a downstream intergenic regulatory site in neurons

In the nucleus, the genome is arranged into self-interacting TADs in which DNA sequences contact each other frequently (Dixon et al., 2012; Nora et al., 2012). Strengthening of intra-TAD and depletion of inter-TAD contacts have been seen during neuronal development, and new TAD boundaries form near developmentally regulated genes as they become transcriptionally activated (Bonev et al., 2017). Analysis of published high resolution HiC data from mouse embryonic stem cells (ESCs) differentiated into NPCs and cortical neurons (CNs) (Bonev et al., 2017) revealed a sub-TAD encompassing the *Bdnf* gene and a downstream gene-free region adjacent to the closest downstream gene, *Lin7c* (**Supplementary Fig. S1A**). The sub-TAD falls at the 3’ end of a large TAD, and its contact frequencies were similar in NPCs and CNs, despite the dramatic difference in *Bdnf* expression. *Lin7c* is expressed in neurons and regulates postsynaptic density (Butz et al., 1998). The expression of *Lin7c* also increased during the differentiation of from NPCs into PMNs (**Fig. 1D**).

CTCF (CCCTC-binding factor) and cohesin are key regulators of TAD boundaries (Nora et al., 2017; Sofueva et al., 2013), and bind to the *Bdnf* locus at promoter IV and intron 7 in mouse primary cortical neurons (Chang et al., 2010). Loss of either CTCF or cohesin compromises *Bdnf* expression from promoter IV following depolarisation, increasing repressive histone modifications (Chang et al., 2010). CTCF binds to *Bdnf* in mouse hippocampus, and loss of CTCF alters intra-gene chromosome looping and reduces *Bdnf* induction in response to fear conditioning (Sams et al., 2016). Chromatin Immunoprecipitation with sequencing (ChIP-seq) analysis identified CTCF peaks in the *Bdnf* and *Lin7c* genes in both NPCs and CNs at sites coinciding with the sub-TAD boundaries (**Supplementary Fig. S1A**). CTCF binding site 1 spans *Bdnf* exon II, but the highest enrichment of CTCF was observed at *Bdnf* binding site 2, located on the downstream part of exon VII and extending into the intron. CTCF binds within *Lin7c* at exon IV. Rad21 ChIP-qPCR at these sites confirmed cohesin binding to *Bdnf* and *Lin7c* CTCF sites in primary NPCs and PMNs (**Supplementary Fig. S1B**). These data indicate that during neuronal development *Bdnf* and *Lin7c* co-occupy a sub-TAD with CTCF-positive, cohesin-positive boundaries.

The chromatin loops which constitute TADs are often cell-type specific and promote accurate gene regulation (Brookes and Riccio, 2019). Sub-TAD level loops often reflect contacts occurring between gene promoters and enhancers, which enable appropriate spatial and temporal control of gene transcription (Fanucchi et al., 2013; Tan-Wong et al., 2012). We reasoned that potential new enhancers of the *Bdnf* gene could be found within the sub-TAD identified here. To identify potential new enhancers of the *Bdnf* gene, we used 4C-seq, a technique that identifies chromatin regions that make contact with a specific ‘viewpoint’ sequence (van de Werken et al., 2012). A viewpoint designed at *Bdnf* exon I identified two regions of interaction in NPCs and PMNs, and in cortical neurons (**Fig. 2A**). The first is internal to the *Bdnf* gene and is located around exon VIII. The second is an intergenic region located downstream (around 170 kb) of *Bdnf* and peaking ∼5 kb upstream of the *Lin7c* gene (Distal Interacting Site, DIS; **Fig. 2A**). The profile was the same irrespective of the *Bdnf* expression levels in the cell type, consistent with the HiC *Bdnf*-*Lin7c* sub-TAD data (**Supplementary Fig. S1A**). To confirm the loop, we prepared a reverse viewpoint from the distal interaction site upstream of *Lin7c* in cortical neurons. This region showed increased contact frequency at sites within the intergenic region, as well as a reciprocal interaction to exon VIII of *Bdnf* (**Supplementary Fig. S2A**).

**Figure 2.**
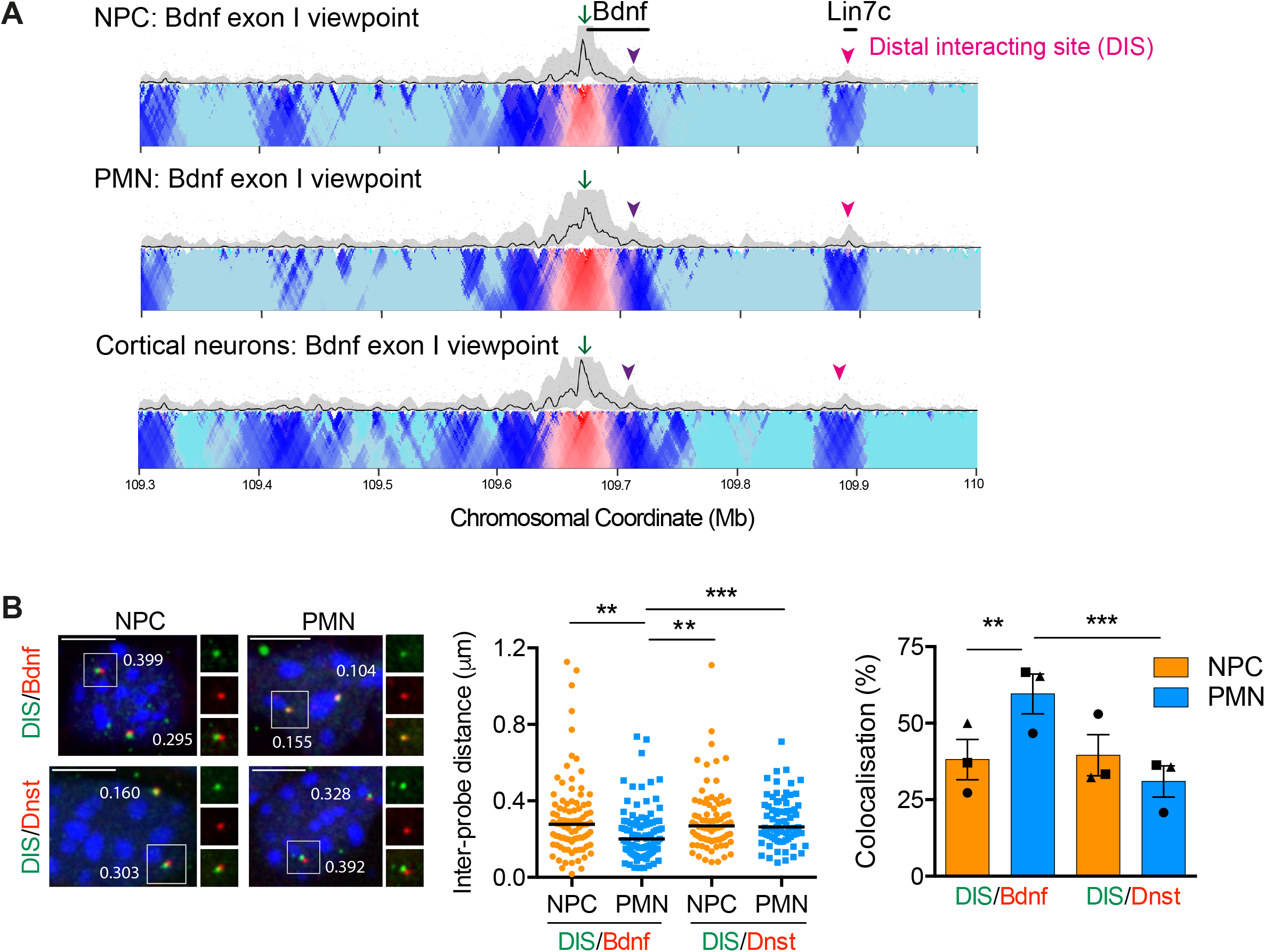
*Bdnf* forms a chromatin loop with an intergenic region and the *Lin7c* gene. See also Supplementary Figure S1 and S2. **A)** Contact profiles from 4C-seq experiments in NPC, PMN and cortical neurons from exon I viewpoint. Interactions to an intragenic site (purple arrowhead) and to a distal interacting site (DIS; pink arrowhead) are indicated. Each image shows a representative 4C-seq experiment (from *n*=2) represented by the median normalized 4C-seq coverage in a sliding window of 51kb (top) and a multi-scale domainogram indicating normalised mean coverage in windows ranging between 2 and 50⍰ kb. **B)** Double DNA-FISH of the enhancer with either the *Bdnf* gene or an equidistant region downstream. Left panel, representative maximal intensity projections of double DNA FISH in NPCs and PMNs. Nuclei were counterstained with DAPI (blue). Scale bar, 5 μm. Middle panel, scatter dot plot of interprobe distance measurements in NPC (orange) and PMN (blue) cells. Solid lines denote medians. \*\**p* < 0.01, \*\*\**p* < 0.001, One-way ANOVA with Dunn’s multiple comparisons (two-tailed). *n* = 87 (DIS/*Bdnf*-NPC), 98 (DIS/*Bdnf*-PMN), 78 (DIS/Dnst-NPC), 74 (DIS/Dnst-PMN) foci across 3 biological replicates. Probe labelling denoted in coloured font. DIS/*Bdnf*-NPC vs. DIS/*Bdnf*-PMN *p*=0.0023; DIS/*Bdnf*-PMN vs. DIS/Dnst-NPC *p*=0.0022; DIS/*Bdnf*-PMN vs. DIS/Dnst-PMN *p*=0.0008. Right panel, colocalisation (defined as an inter-probe distance of 225nm or less) of FISH signals in double DNA FISH experiments performed in NPCs and PMNs. Bars represent mean ± SEM, and points show results from different biological replicates (*n* = 3). ** *p* < 0.01, *** *p* < 0.001, Fisher’s exact test (two-tailed). DIS/*Bdnf*-NPC vs. DIS/*Bdnf*-PMN p=0.0051; DIS/*Bdnf*-PMN vs. DIS/Dnst-PMN p=0.0002.

To investigate cell-to-cell variation in the loop, and study whether it changed during neuronal differentiation, we employed a single cell assay. DNA-FISH was performed using fosmids encompassing a) the DIS and *Lin7c* gene, b) the *Bdnf* gene, and c) a downstream region located at the same distance from the DIS as *Bdnf* (169 kb). The distance between these probes analysed in pairwise sets revealed that in PMNs the putative enhancer was closer to, and exhibited more frequent interactions with, the *Bdnf* probe, compared to the downstream probe (**Fig. 2B**). This confirms the sub-TAD of increased interaction encompassing the *Bdnf* gene and the DIS, compared to downstream regions, that we identified using 4C-seq (**Fig. 2A**). The DIS and *Bdnf* probes were found in closer proximity in PMN than in NPCs, and the colocalisation frequency increased during differentiation (**Fig. 2B**). Thus, although the looping profiles are similar at the population level (**Fig. 2A**), single cell analysis indicated that an increase in interaction takes place during neuronal differentiation (**Fig. 2B**). The use of a reciprocal combination of labels on the probes supported this conclusion (**Supplementary Fig. S2B**). This is in accordance with previous studies showing that detection of interaction in non-expressing cells using chromosome conformation capture technologies may reflect proximity of the enhancer and promoter, while the increased colocalisation seen with DNA-FISH demonstrates direct interactions upon induction (Fudenberg and Imakaev, 2017; Williamson et al., 2016).

### The distal interacting site bears many characteristics typical of enhancers

To assess whether the intergenic region contacting *Bdnf* exhibits the characteristics of an enhancer, we first analysed publicly available data. Sensitivity to DNase I is a feature of active chromatin regions including promoters and enhancers (Boyle et al., 2008). ENCODE DNase I hypersensitivity data showed peaks in the intergenic region in whole brain (**Fig. 3A**). DNase I hypersensitivity at the DIS showed a similar pattern across tissues to that observed at *Bdnf* promoters and was found only in embryonic and adult brain (**Supplementary Fig. S3A**). *Lin7c* showed a different pattern, with DNase I hypersensitivity also detected at the transcription start site (TSS) in other tissues, including lung and liver (**Supplementary Fig. S3A**), suggesting that the DIS is not simply part of the *Lin7c* promoter region. A dataset using an alternative chromatin accessibility assay named Assay for Transposase-Accessible Chromatin with Sequencing (ATAC-seq) (Su et al., 2017), also identified open chromatin at the putative *Bdnf* enhancer in microdissected hippocampal dentate gyri (not shown).

**Figure 3.**
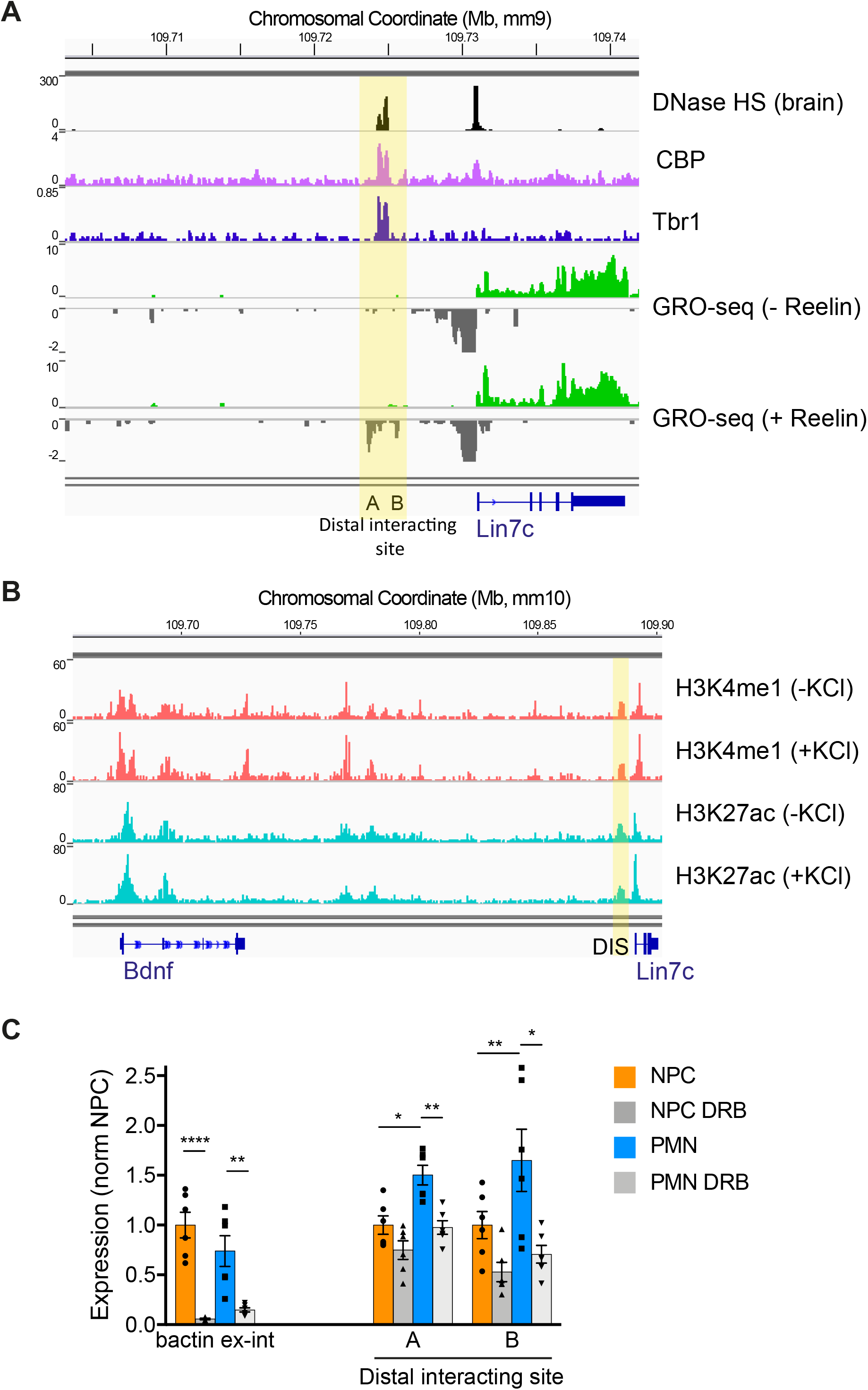
The *Bdnf*-interacting intergenic region displays hallmarks of an enhancer. See also Supplementary Figure S3. **A)** Available data for DNaseI hypersensitivity (ENCODE), and CBP (Telese et al., 2015) and Tbr1 (Notwell et al., 2016) ChIP-seq were visualised at the distal interacting site in brain or neurons to assess chromatin accessibility and transcription factor binding. GRO-seq profiles from control or Reelin-treated cortical neurons (Telese et al., 2015) show a peak of nascent transcription at the putative *Bdnf* enhancer in stimulated cortical neurons. A and B mark sites positive for GRO-seq signal within the putative enhancer region that were used for qRT-PCR verification. **B)** H3K4me1 (pink) and H3K27ac (blue) ChIP-seq (Policarpi et al., 2017) in cortical neurons minus (-) and plus (+) KCl show enhancer markers at the putative *Bdnf* enhancer. **C)** qRT-PCR expression analysis of two regions within the putative enhancer in untreated or DRB-treated NPC and PMN. Data are normalised to NPC. ß-actin primary transcripts shown to demonstrate DRB treatment efficacy. Bars represent mean ± SEM, and points show results from different biological replicates (*n* = 6). ^*^p < 0.05, *^**^p < 0.01, ^***^p < 0.001, ^****^p < 0.0001, two-way ANOVA with Sidak’s multiple comparisons test. ß-actin ex-int: p<0.0001 (NPC Unt vs. DRB), p=0.0011 (PMN Unt vs. DRB). Enh-A: p=0.0009 (PMN Unt vs. DRB), p=0.0014 (Unt NPC vs. PMN). Enh-B: p=0.0032 (PMN Unt vs. DRB), p=0.0410 (Unt NPC vs. PMN).

We then investigated other enhancer hallmarks using publicly available ChIP-seq datasets (Notwell et al., 2016; Policarpi et al., 2017; Telese et al., 2015) (**Supplementary Table 1**). Chromatin modifications, in particular H3K4me1 and H3K27ac, are known to predict enhancer function genome-wide (Creyghton et al., 2010; Heintzman et al., 2007). The histone acetyltransferase CBP (CREB Binding Protein) is a known enhancer regulator, which catalyses the addition of H3K27ac (Rada-Iglesias et al., 2011; Tie et al., 2009). The transcriptional coactivator Mediator interacts with cohesin to regulate enhancer-promoter looping (Kagey et al., 2010). Enhancers are also sites of multiple transcription factor recruitment (Carullo and Day, 2019). We identified a clear peak of the enhancer chromatin markers H3K27ac and H3K4me1 at the DIS in both basal and depolarised cortical neurons (**Fig. 3B**). CBP and Mediator were also found to bind to the putative *Bdnf* enhancer (**Fig. 3A, Supplementary Fig. S3B**) as well as the transcription factors Mef2, Creb and Tbr1 (**Fig. 3A, Supplementary Fig. S3B**), which regulates neuronal chromatin folding (Bonev et al., 2017). The transcription factors and coactivators show a double peak at the intergenic region, coinciding with a double peak of DNase I hypersensitive sites.

Enhancers can be transcribed in many cell types, including neurons (Kim et al., 2010; Policarpi et al., 2017; Telese et al., 2015). In some instances, the enhancer RNA (eRNA) has functional roles, such as interacting with NELF (Negative Elongation Factor) (Schaukowitch et al., 2014), CBP (Bose et al., 2017), or RNAPII (Policarpi et al., 2017), or affecting 3D contacts (Li et al., 2013). In other systems, enhancer RNA transcription may contribute to the maintenance of the transcriptional machinery or the opening of the chromatin (Mousavi et al., 2013; Panigrahi et al., 2018). Regardless of mechanism, the production of eRNAs is now considered a critical feature of active enhancers. We therefore sought to determine whether transcriptional activity could be detected from the putative *Bdnf* enhancer. eRNAs are lowly expressed and unstable and, in keeping with this, conventional RNA-seq databases do not always show RNA transcription at enhancer sites. Methods that detect nascent RNA such as Genome Run On with sequencing (GRO-seq) are better suited for detecting eRNAs as they map transcriptionally engaged RNAPII genome-wide (Core et al., 2008). Analysis of GRO-seq data from Reelin-stimulated cortical neurons (Telese et al., 2015), showed that RNA is transcribed from the negative strand of the DIS region (**Fig. 3A**). GRO-seq did not detect bidirectional transcriptional activity, which may be due to low expression levels of the eRNA and the unidirectional nature of eRNA transcription at the single cell level (Kouno et al., 2019); however the close-by *Lin7c* gene exhibited divergent RNA production at the active promoter, as expected (Core et al., 2008; Seila et al., 2008).

To validate the sequencing data, qRT-PCR was performed on NPCs and PMNs using primers that generate amplicons within the region of GRO-seq enrichment (**Fig. 3A**, sites A and B). Since eRNA are transcribed at very low levels, cells were treated with the transcriptional inhibitor DRB (5,6-dichloro-1-beta-D-ribofuranosylbenzimidazole) to determine the background level. We found that the putative enhancer region was transcribed in PMNs, at levels significantly higher than either in NPCs or in DRB-treated PMNs (**Fig. 3C**). Taken together, these finding demonstrate that the intergenic region interacting with *Bdnf* possesses the hallmarks of an enhancer and is transcribed in PMNs.

### The distal interacting site is a *Bdnf* enhancer that regulates *Bdnf* expression during neuronal differentiation

To test whether the intergenic region is a functional enhancer for *Bdnf* during NPC differentiation we employed RNA-guided Clustered Regularly Interspaced Palindromic Repeats inhibition (CRISPRi). A catalytic mutant Cas9 (dCas9) was fused to a transcriptional inhibitor (dCas9-KRAB) (Thakore et al., 2015) and lentivirus were generated, either in combination with no targeting gRNA (Empty) or targeted to the putative enhancer region (Enh^g1^, Enh^g2^). NPCs were infected with CRISPRi lentivirus and allowed to differentiate *in vitro*. Immunofluorescence confirmed efficient targeting of PMNs at DIV 7 (**Fig. 4A**; >60% GFP+). Interestingly, compared to untargeted CRISPRi, inhibition of the enhancer region caused a dispersion of the PMN clusters derived from the NPC rosettes, quantified as an increased nuclei-nuclei distance (**Fig. 4A, B**). Neuronal dispersion was reversed by inclusion of lentiviral expressed *Bdnf* (**Fig. 4C**), suggesting that *Bdnf* is necessary for neuron-neuron interaction and the formation of neuronal clusters *in vitro*. The expression of markers of neuronal differentiation (Map2, NeuN, *Nestin*) did not change (not shown), suggesting that inhibition of *Bdnf* expression may specifically influence cell migration or adhesion properties.

**Figure 4.**
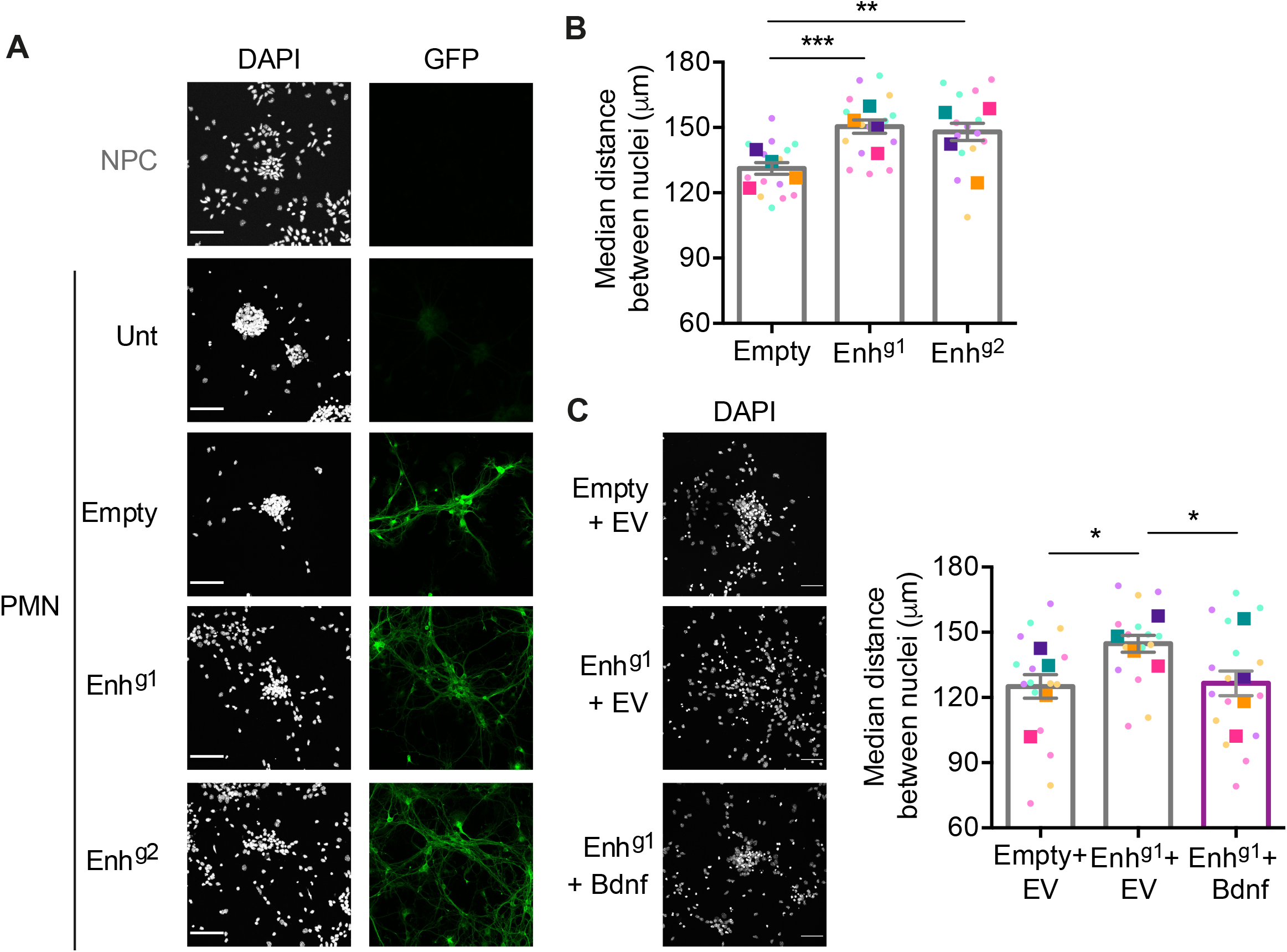
Transcription from the putative enhancer is required for neuronal clustering. **A)** Representative maximal intensity projections of NPC and PMN, and PMN treated with lentiviral CRISPRi (Empty or targeting the putative enhancer region (Enh^g1^, Enh^g2^)). DAPI images (grey) show dispersion of cells within clusters; GFP staining (green) shows percentage of lentiviral-targeted cells. Scale bar, 50 μm. **B)** The median distance between cells per image (image taken to include a single cluster and any surrounding cells) is increased in PMNs when *Bdnf* enhancer function is compromised. Small dot points show values from individual images colour-coded according to the biological replicate; large square points show means of each biological replicate (*n*=14 over 4 biological replicates (Empty), 15 over 4 biological replicates (Enh^g1^), 14 over 4 biological replicates (Enh^g2^)). Bars represent means ± SEM. \**p* < 0.05, \*\**p* < 0.01, \*\*\**p* < 0.001, unpaired t test (two-tailed). Empty vs. Enh^g1^ *p*=0.0008, *t*=3.764, *df*=27; Empty vs. Enh^g2^ *p*=0.0057, *t*=3.012, *df*=26. **C)** Expression of *Bdnf* rescues the increase of cell spacing following enhancer inhibition. Quantification of the median distance between cells per image (image taken to include a single cluster and any surrounding cells) for CRISPRi experiments including control (EV; Empty vector) or *Bdnf*-expressing lentivirus. Light points show values from individual images colour-coded according to the biological replicate they belong to; bright points show means of each biological replicate (*n*=16 over 4 biological replicates (Empty+EV), 15 over 4 biological replicates (Enh^g1^+EV), 16 over 4 biological replicates (Enh^g1^+*Bdnf*)). Bars represent means ± SEM. *p < 0.05, unpaired t test (two-tailed). Empty+EV vs. Enh^g1^+EV *p*=0.0251, *t*=2.363, *df*=29; Empty+EV vs. Enh^g1^+*Bdnf p*=0.0387, *t*=2.166, *df*=29.

To analyse the effect of enhancer inhibition on specific *Bdnf* isoforms, we generated CRISPRi lentiviral particles incorporating a puromycin resistance cassette and selected transduced neurons for two days prior to harvesting the PMNs. Induction of eRNA during differentiation was significantly decreased in the presence of enhancer-targeted guide RNAs (**Fig. 5A**). Enhancer inhibition caused a significant reduction of *Bdnf* mRNA (measured in the universal exon) confirming that that we have identified a functional *Bdnf* enhancer (**Fig. 5B**). Analysis of different *Bdnf* isoforms indicated that eRNA inhibition resulted in significant reduction of *Bdnf* exon I, IV, VI, VIII and IXa variant transcription (**Fig. 5B**). We did not detect significant changes in *Bdnf* exon II or V variants (**Fig. 5C**). Importantly we did not see a reduction in *Lin7c* expression (**Fig. 5C**), confirming that the CRISRPi inhibitory effect at the enhancer does not spread into the *Lin7c* promoter.

**Figure 5.**
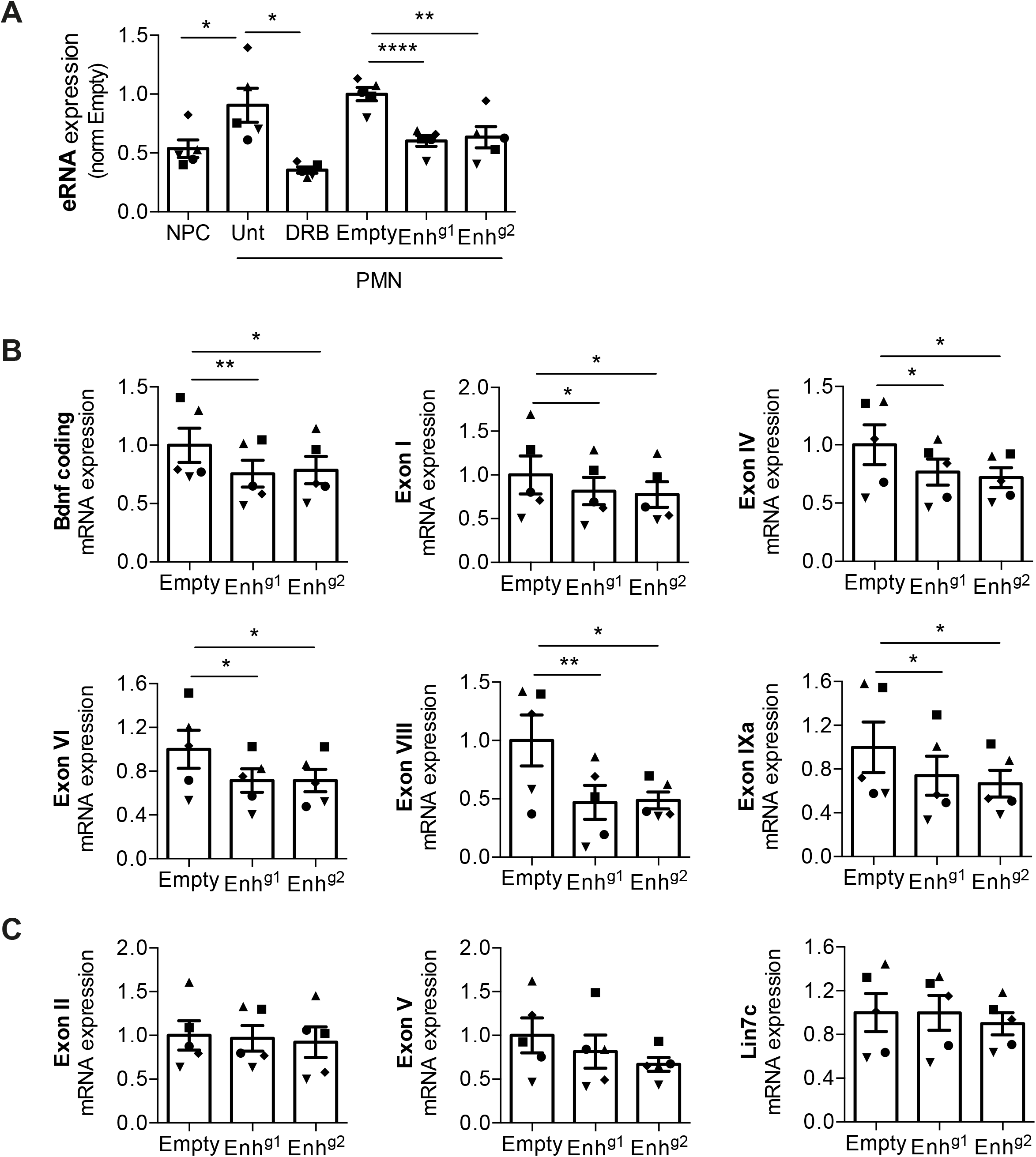
The enhancer is required for full *Bdnf* induction during differentiation. Expression profiles of **A)** the enhancer RNA and **B)** the *Bdnf* transcript variants and coding isoform that are insensitive to enhancer inhibition. qRT-PCR of PMN targeted with lentiviral dCas9-KRAB targeted by no guide (Empty) or guides against the enhancer (Enh^g1^ and Enh^g2^). Data are normalised to Empty vector transduced cells. Bars represent mean ± SEM, and points show different biological replicates (*n* = 5). \**p* < 0.05, \*\**p* < 0.01, \*\*\**p* < 0.001; paired *t*-test (two-tailed). eRNA: NPC vs. PMN-Unt *p*=0.0105, *t*=4.539, *df*=4; PMN-Unt vs. PMN-DRB *p*=0.0151, *t*=4.077, *df*=4; PMN-Enh^g1^ vs. PMN-Empty p<0.0001, t=19.74, *df*=4; PMN-Enh^g2^ vs. PMN-Empty p=0.0014, t=7.954, *df*=4. *Bdnf* coding: Empty vs. Enh^g1^ *p*=0.0036, *t*=6.108, *df*=4; Empty vs. Enh^g2^ *p*=0.0246, t=3.512, *df*=4. Exon I Empty vs. Enh^g1^ p=0.0411, *t*=2.972, *df*=4; Empty vs. Enh^g2^ *p*=0.0375, *t*=3.065, *df*=4. Exon IV Empty vs. Enh^g1^ *p*=0.0210, *t*=3.694, *df*=4; Empty vs. Enh^g2^ *p*=0.0315, *t*=3.247, *df*=4. Exon VI Empty vs. Enh^g1^ p=0.0145, t=4.134, *df*=4; Empty vs. Enh^g2^ *p*=0.0229, *t*=3.592, *df*=4. Exon VIII Empty vs. Enh^g1^ *p*=0.0091, *t*=4.729, *df*=4; Empty vs. Enh^g2^ *p*=0.0417, *t*=2.956, *df*=4. Exon IXa Empty vs. Enh^g1^ *p*=0.0363, *t*=3.099, *df*=4; Empty vs. Enh^g2^ *p*=0.0475, *t*=2.826, *df*=4. **C)** Expression profile by qRT-PCR of the *Bdnf* transcript variants and of the *Lin7c* mRNA that are insensitive to enhancer inhibition. Data are normalised to Empty vector transduced cells. Bars represent mean ± SEM, and points show different biological replicates (*n* = 5).

### The *Bdnf* enhancer regulates dendritogenesis in cortical neurons

*Bdnf* expression is necessary for activity-dependent dendritogenesis (McAllister et al., 1996; McAllister et al., 1995), a process critical for neuronal growth at later stages of development. To assess whether the *Bdnf* enhancer played a role in these processes we first investigated whether it is transcribed in an activity-dependent manner. E15.5 cortical neurons were stimulated with KCl and eRNA levels were assessed with qRT-PCR. *Bdnf* eRNA was significantly increased concomitant with an increase in *Bdnf* gene expression (**Fig. 6A**) and the activity-dependent complex AP-1 was recruited in response to neuronal depolarisation (**Supplementary Fig. S4A**). This is consistent with transcription factors encoded by early response genes, like Fos and Jun, controlling the expression of late response genes, such as *Bdnf* (Malik et al., 2014).

**Figure 6.**
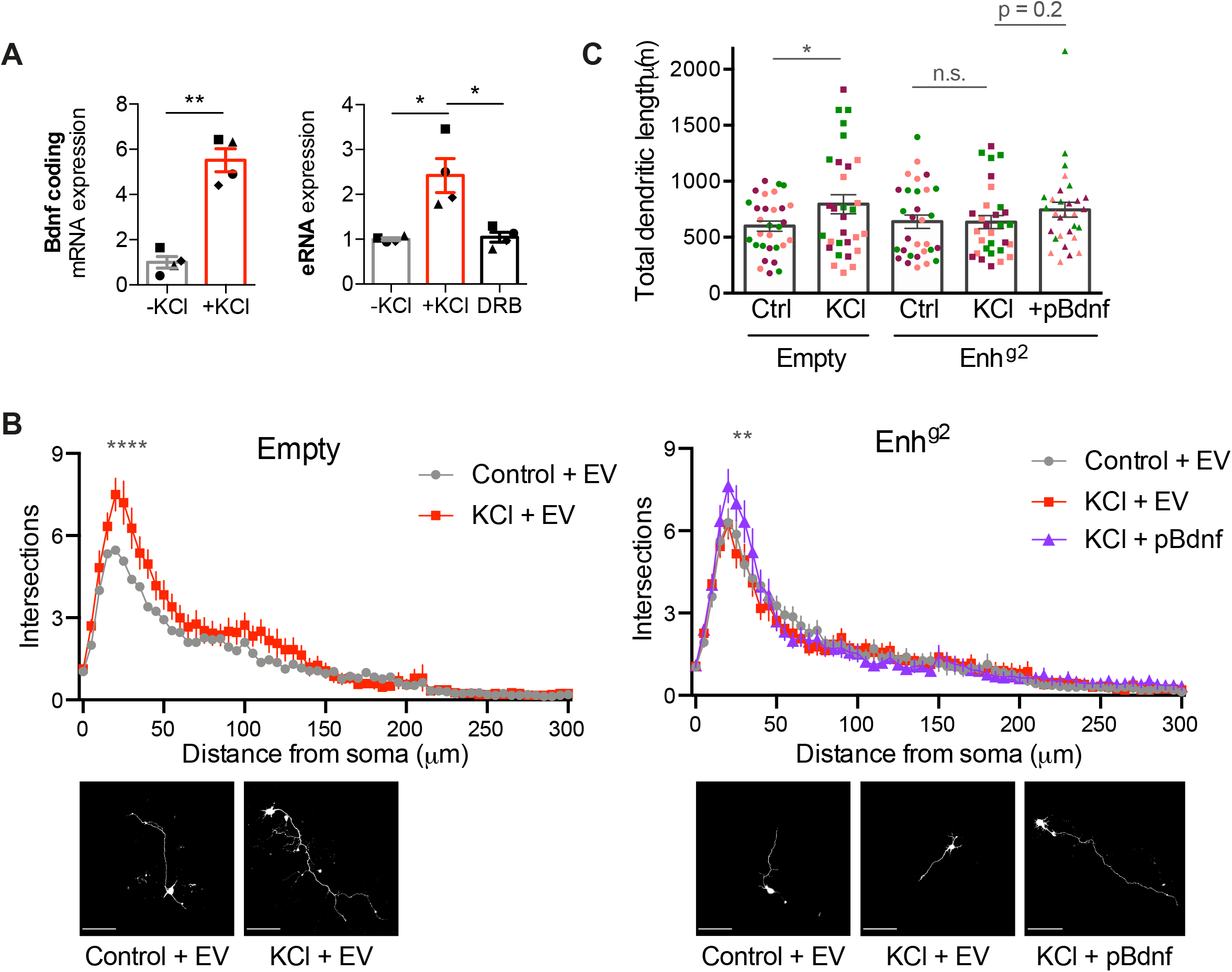
The enhancer is required for activity-dependent dendritogenesis. See also Supplementary Figure S4. **A)** Enhancer RNA and *Bdnf* mRNA expression increase following neuronal depolarisation. Expression profile of the enhancer RNA and *Bdnf* coding mRNA in cortical neurons maintained in basal (-KCl Control) or depolarising (50 mM +KCl) conditions for 48h. Levels assessed by qRT-PCR and normalized to -KCl samples. Bars represent mean ± SEM, and points show values of different biological replicates (*n* = 4). *p* < 0.05, *p* < 0.01; paired t-test (two-tailed). *Bdnf* coding -KCl vs. +KCl *p*=0.0020, *t*=10.27 *df*=3. eRNA -KCl vs. +KCl *p*=0.0338, *t*=3.721 *df*=3, +KCl vs. DRB *p*=0.0161, *t*=4.924 *df*=3. **B)** Sholl analysis of the dendritic processes of 30 neurons per condition (10 per biological replicate). For each distance point, the mean number of intersections ± SEM is shown. \*\**p* < 0.01, \*\*\**p* < 0.001, \*\*\*\**p* < 0.0001, two-way ANOVA with Sidak’s multiple comparisons test. Empty: Control+EV vs. KCl+EV 20 μm from soma p<0.0001, mean diff -2.033, 95% CI of diff - 3.233 to -0.8332; 25 μm from soma *p*<0.0001, mean diff -2.133, 95% CI of diff -3.333 to - 0.9332; 30 μm from soma p<0.0001, mean diff -1.867, 95% CI of diff -3.067 to -0.6665; 35 μm from soma p=0.0351, mean diff -1.233, 95% CI of diff -2.433 to -0.03321; 40 μm from soma *p*=0.0006, mean diff -1.567, 95% CI of diff -2.767 to -0.3665. Enh^g2^: KCl+EV vs. KCl+p*Bdnf* 20 μm from soma *p*=0.0022, mean diff -1.400, 95% CI of diff -2.546 to -0.2538; 25 μm from soma *p*<0.0001, mean diff -1.833, 95% CI of diff -2.980 to -0.6871; 30 μm from soma *p*=0.0022, mean diff -1.400, 95% CI of diff -2.546 to -0.2538. Lower panel, representative images of cortical neurons transfected with a pCIG GFP expression vector (EV or p*Bdnf*) in combination with dCas9-KRAB-MECP2 and an expression vector for guide RNAs (Empty or targeting the putative enhancer region (Enh^g2^)). Cells were maintained under basal or depolarising (KCl, 50 mM) conditions for 48 hr, followed by GFP immunostaining. Scale bar, 100 μm. **C)** Quantification of the total length of the dendritic processes of neurons analysed in (b). Bars show mean ± SEM; points show each data for each neuron coloured by biological replicate (*n*=3). \**p* < 0.01, unpaired t-test (two-tailed). Empty: Control+EV vs. KCl+EV *p*=0.0489, *t*=2.011, *df*=58. Enh^g2^: KCl+EV vs. KCl+p*Bdnf p*=0.2095, t=1.269, *df*=58.

To analyse the role of the *Bdnf* enhancer in activity-dependent dendritogenesis, cortical neurons were transfected with plasmids encoding the potent repressor dCas9-KRAB-MECP2 (Yeo et al., 2018), along with gRNAs (BPK1520: Empty, Enh^g1^, Enh^g2^) and GFP. Neurons were maintained in basal or depolarising conditions (50 mM KCl) for 48 hours, and dendrites from GFP-positive, non-overlapping neurons were traced. As expected, depolarisation induced a significant increase in dendritic length and complexity in control neurons transfected with dCas9-KRAB-MECP2 only (Empty, **Fig. 6B, C; Supplementary Fig. S4B-D**). In contrast, KCl-dependent dendritic growth or branching was abolished when neurons were transfected with dCas9-KRAB-MECP2 targeted to the putative enhancer (Enh^g1^ or Enh^g2^, **Fig. 6B, C; Supplementary Fig. S4B-D**). To assess whether the effect of enhancer inhibition depended on *Bdnf* gene expression, dendritogenesis was assessed in neurons expressing a vector encoding the *Bdnf* coding sequence (p*Bdnf*) or an empty control vector (EV), and co-transfected with CRISPRi vectors (Empty or Enh^g2^). Depolarisation of control cortical neurons increased dendritic growth and arborisation, which was reduced by enhancer inhibition (**Fig. 6B, C**). Co-transfection of the p*Bdnf* plasmid rescued the defect in branching close to the soma, although it did not fully reinstate the total length or the branching in distal dendrites (**Fig. 6B, C**). Together these data indicate that the newly identified *Bdnf* enhancer regulates *Bdnf* expression to promote activity-dependent dendritic growth.

## Discussion

Elucidating how the *Bdnf* gene is regulated is paramount to a complete understanding of its critical neurotrophic actions during development and in mature neurons. Here, we identify a novel enhancer that influences *Bdnf* expression during neuronal differentiation and in response to neuronal depolarisation. We demonstrated that, in addition to displaying most enhancer hallmarks, including binding of CBP and Mediator (**Fig. 3A, Supplementary Fig. S3B**), chromatin accessibility (**Fig. 3A, Supplementary Fig. S3A**), histone modifications (**Fig. 3B**) and transcription (**Fig. 3A, C**), enhancer activity was required for full activation of the *Bdnf* gene during neuronal progenitor differentiation (**Fig. 5**). Abrogation of enhancer function altered cellular interactions (**Fig. 4**) and activity-dependent dendritic growth (**Fig. 6, Supplementary Fig. S4**). The enhancer loops to the *Bdnf* gene in neurons (**Fig. 2, Supplementary Fig. S2**), and analysis of genome topology revealed that the gene and enhancer are located within a sub-TAD which is bounded by CTCF and cohesin (**Supplementary Fig. S1A, B**). *Bdnf* activation correlates with increasing frequency of enhancer-promoter co-localisation (**Fig. 2B, Supplementary Fig. S2B**) and movement of the genomic region away from the nuclear periphery (**Fig. 1E**).

### A novel enhancer that regulates *Bdnf* expression during neuronal differentiation

We discovered an enhancer that regulates nearly all variants of *Bdnf*, albeit to different extents, in differentiating neurons (**Fig. 5B**). In keeping with this, we observed a significant effect of enhancer inhibition on total *Bdnf* mRNA expression (**Fig. 5B**). Interestingly, this effect was not seen upon inhibition of the intronic enhancer (Tuvikene et al., 2021). The intronic enhancer regulates transcription from *Bdnf* promoters I, II, and III, but not later promoters (Tuvikene et al., 2021), therefore showing a very different specificity to the enhancer identified here and suggesting that multiple regulatory regions are responsible for spatially and temporally restricted *Bdnf* expression. *Bdnf* transcript isoforms show different expression patterns across brain regions, cell types, developmental stages and stimuli, and further investigations will be needed to address how each *Bdnf* enhancer contributes to this. For example, during differentiation, we did not observe an effect of the intergenic enhancer on exon II-containing mRNA (**Fig. 5C**). An REI element, bound by the REST repressor (Timmusk et al., 1999), has been described within exon II, and it is possible that this element insulates exon II from the effect of the enhancer (Tang et al., 2021).

Importantly, we also demonstrate that enhancer inhibition has significant physiological consequences. Inhibition of the novel enhancer causes a dispersion of neuronal clusters (**Fig. 4**) and prevents activity-dependent dendritogenesis (**Fig. 6, Supplementary Fig. S4**). Both cellular consequences are mitigated by *Bdnf* expression (**Fig. 4C, Fig. 6B, C**). Enhancer inhibition abrogated activity-induced dendritic growth, but inclusion of p*Bdnf* only rescued this defect close to the soma (**Fig. 6B**). Comparing p*Bdnf*-transfected neurons with EV-transfected cells in basal conditions (-KCl Control) showed an increase in dendritic branching close to the soma, but not more distally (not shown), suggesting that this is where *Bdnf* exerts its greatest effect. Hence, the additional loss of distal branching after enhancer inhibition (**Fig. 6B**) may be due to secondary or nonspecific effects. The effect of the *Bdnf* enhancer on dendritic growth is likely due to an autocrine mechanism as very few neurons are transfected in this assay. However, the effect on neuronal clustering may be either autocrine, paracrine or both, since the majority of cells in the population were affected.

### Complex genome topology around the *Bdnf* locus

We explored the 3D genome architecture of the *Bdnf* genomic region using HiC and 4C-seq data and described for the first time a sub-TAD of increased interaction frequency that includes the *Bdnf* gene and the downstream intergenic region into *Lin7c*, with CTCF and cohesin-positive boundaries within *Bdnf* and *Lin7c* genes (**Fig. 2A, Supplementary Fig. S1A, B, S2A**). Within this sub-TAD we identified a chromatin loop linking a distal intergenic region to the *Bdnf* gene, and an intragenic loop from *Bdnf* exon I to exon VIII (**Fig. 2A, Supplementary Fig. S2A**), suggesting that the enhancer-promoter loop may be anchored around exon VIII. Neither the sub-TAD boundaries nor the enhancer-promoter loop sites changed during neuronal differentiation (**Fig. 2A, Supplementary Fig. S1A**), suggesting that in NPCs they are prewired. Topological structure has been shown to precede gene activation in a number of studies (Jin et al., 2013; Kolovos et al., 2016; Montavon et al., 2011; Paliou et al., 2019; Rubin et al., 2017), and it has been suggested that preconfigured loops prime genes for transcriptional induction (de Laat and Duboule, 2013). However, single cell imaging showed an increase in colocalisation of enhancer with promoter from NPC to PMN (**Fig. 2B, Supplementary Fig. S2B**). This is consistent with the 4C-seq and HiC data reflecting proximity of the sequences, while the DNA-FISH detects interaction between sites in cell types where the gene is expressed (Williamson et al., 2016). This demonstrates the added value of complementing chromosome conformation capture data with imaging techniques, and suggests that the sub-TAD organisation may facilitate the interaction of the distal enhancer with *Bdnf*.

A recent study used 5C-seq to examine the topology of the *Bdnf* genomic region in cortical neurons (Beagan et al., 2020). Importantly, the enhancer-promoter loop that we identified with 4C is also found in data generated in this study on basal and stimulated neurons. In addition to constitutive loops, the 5C-seq also describes novel looping events which form in response to depolarization (Beagan et al., 2020). Future investigations will clarify the functional significance of these loops on *Bdnf* expression, and the interplay with the enhancer characterised here.

### The *Bdnf* enhancer is conserved in human cells

Understanding the different facets of *BDNF* regulation has important implications for a variety of physiological processes and for the pathogenesis of neurological disorders in which *BDNF* is downregulated. The enhancer sequence that we identified in mouse is conserved in the human genome, where it is located in a similar orientation and position relative to the *BDNF* and *LIN7C* genes. In humans, an antisense transcript that regulates *BDNF* expression, *BDNF*-*AS*, runs from immediately upstream of the *LIN7C* TSS through the intergenic region and the *BDNF* gene itself (Modarresi et al., 2012; Pruunsild et al., 2007). Integration of enhancer function with *BDNF*-*AS* transcription will need to be considered.

In conclusion, we have identified a novel enhancer for the neurotrophin-encoding *Bdnf* gene. Enhancer activity is required for appropriate upregulation of many *Bdnf* transcript variants, as well as total *Bdnf* mRNA levels during neuronal differentiation. Moreover, enhancer inhibition alters neuronal clustering during differentiation and abrogates activity-dependent dendritic growth, suggesting that the eRNA regulates *Bdnf* function in diverse physiological contexts. Different *Bdnf* transcript isoforms display different spatiotemporal expression patterns, and also show specific changes in neurological disease. Here, we add an important piece to the puzzle of understanding the regulation of this essential growth factor gene, which may be important in understanding its dysregulation in pathological states.

## Acknowledgements

We thank Catia Andreassi (Riccio Lab) for constant advice and helpful discussion, and Janos Kriston-Vizi (UCL) for advice on image analysis. We thank Dimitra Georgopoulou (UCL) for 4C-seq advice. We thank Tommaso Squeri (Kings College London) and Charlotte Laurent (Imperial College London) for their work on this project during their summer placements. We thank all members of the Riccio laboratory for valuable input and discussions. This project has received funding from the European Union’s Horizon 2020 research and innovation programme under the Marie Skłodowska-Curie grant agreement Number GA702327 (to E.B.). This work was supported by the Wellcome Trust Investigator Awards 103717/Z/14/Z and 217213/Z/19/Z (to A.R.) and 106985/Z/15/Z (to S.H.), and the MRC LMCB Core Grant MC/U12266B (to A.R).

## Author Contributions

E.B. performed most experiments and helped conceive the project and write the manuscript. H.Y.A.A. performed the experiments in Supplementary Fig. 4. W.V. analysed HiC and CTCF ChIP-seq data (Supplementary Fig. 1A). C.B. set up analysis of 4C-seq data (Fig. 2A, Supplementary Fig. S2A). S.H. advised on genome topology techniques and interpretation, supervised W.V. and C.B. and contributed to writing the manuscript. A.R. conceived the project and wrote the manuscript.

## Methods

### Cortical progenitor cell culture

All experiments performed in this study were approved by the UK Home Office and were performed under the project license 7813074 held by AR. All animal studies were approved by the Institutional Animal Care and Use Committees at University College London. Cortical progenitor culture was performed essentially as described in (Nitarska et al., 2016). Cortices were dissected from E12.5 C57BL/6J mouse embryos in dissection buffer (2.5 mM Hepes pH 7.4, 30 mM glucose, 1 mM CaCl_2_, 1 mM MgSO_4_, 4 mM NaHCO_3_, 1X HBSS) supplemented with 1 U/ml Dispase I (Sigma) and 0.6 mg/ml DNase I (Sigma). Dissected cortices were digested in dissociation media (1 mM Hepes pH 7.4, 20mM glucose, 98 mM Na_2_SO_4_, 30 mM K_2_SO_4_, 5.8 mM MgCl_2_, 0.25 mM CaCl_2_, 0.001% Phenol red) supplemented with 20 U/ml of papain (Worthington) for 25 min at 37°C. After digestion, cortices were washed, dissociated and plated on Nunc dishes (Thermo Fisher Scientific) or glass coverslips coated with 40 μg/ml poly-D-lysine (Sigma) and 2 μg/ml Laminin (BD Bioscience) in DMEM/F12 medium supplemented with 1X B27, 1X N2, 1 mM glutamine, 1 mM NaHCO_3_ and 10 ng/ml of bFGF (Thermo Fisher Scientific). Cells were plated more densely for NPC cultures harvested after 2 days *in vitro* (DIV) than for PMN cultures harvested at 7 DIV (90 mm dish for 4C-seq and ChIP: NPC 2.5 × 10^6^ cells, PMN 1 × 10^6^ cells; 6-well plates for qRT-PCR analysis: NPC 3.4 × 10^5^ cells, PMN 1.7 × 10^5^ cells; 24-well plates with glass coverslips for imaging: NPC 5.0 × 10^4^ cells, PMN 2.5 × 10^4^ cells). For PMN cultures, after 2 DIV half of the medium was replaced with Neurobasal medium supplemented with 1X B27, 1 mM glutamine and 200 ng/ml NT3 (Alomone labs). After 5 DIV, half of the medium was replaced with Neurobasal medium supplemented with 1X B27, 1 mM glutamine, 200 ng/ml NT3 (Alomone labs) and 20 μM 5-Fluoro-21-deoxyuridine (FdU; Merck). Cells were maintained in 37°C, 5% CO_2_ incubators.

### Cortical neuron culture

Cortical neurons were dissected from E15.5 C57BL/6J mouse embryos and dissociated as above. Neurons were cultured on Nunc dishes (Thermo Fisher Scientific) or glass coverslips coated with 40 μg/ml poly-D-lysine (Sigma) and 2 μg/ml Laminin (BD Bioscience) and plated in MEM supplemented with 10% fetal bovine serum, 5% horse serum, 1 mM glutamine and 1X penicillin-streptomycin. After 2-6 hours, culture medium was replaced with Neurobasal medium supplemented with 1X B27, 1 mM glutamine, 1X penicillin-streptomycin and 10 μM fluorodeoxyuridine (Merck). Cells were cultured at 37°C, 5% CO_2_ for 2-7 days, and one day before the experiment, 2/3 of the plating medium was replaced with medium lacking B27.

### RNA isolation and reverse transcription

For transcriptional inhibition, 50 μM of the RNAPII inhibitor DRB (Merck) was added to culture medium for 1h. RNA was isolated from neuronal cultures using TRIzol (Thermo Fisher Scientific) according to the manufacturer’s instruction. RNA was treated with the TURBO DNA-free kit (Thermo Fisher Scientific) before being reversed-transcribed in a 20 μl reaction volume containing random hexamers, RiboLock RNAse inhibitor and RevertAid (Thermo Fisher Scientific). RT-qPCR reactions (20 μl) contained 12.5 μl SYBR Select Master Mix (Thermo Fisher Scientific) and 0.25 μM (primer sequences shown in **Supplementary Table S3**) and were performed on a BioRad CFX qPCR machine.

### DNA Fluorescence In Situ Hybridization (FISH)

DNA-FISH experiments were performed as described previously (Policarpi et al., 2017) with some modifications. Cells were fixed for 10 min in 4% PFA (TAAB) in PBS, followed by permeabilization for 10 min in 0.5% Triton-X 100 in PBS. After blocking with PBS+ (PBS plus 0.1% casein, 1% BSA, 0.2% fish skin gelatin) for 1h, coverslips were incubated overnight with primary antibodies in PBS+ if necessary. For immuno detection, coverslips were washed in PBS, incubated with appropriate secondary antibodies for 1h in PBS+, and washed in PBS. For DNA-FISH without immunostaining, after the PBS+ block, coverslips were washed in PBS and then proceeded directly to post-fixation. Post-fixation in 4% PFA (TAAB) in PBS (10 min) was followed by permeabilization in 0.1M HCl, 0.7% Triton-X 100 (10 min, on ice), and by denaturation with 70% formamide in 2X SSC (80°C, 30 min). FISH hybridization with probes was carried out overnight at 42°C. Probes (BAC *Bdnf* RP24-149F11 for lamina association; fosmid probes for double DNA-FISH (Enhancer WIBR1-0557J07, *Bdnf* WIBR1-0841J20, Downstream WIBR1-0166C24), BACPAC Resources) were labelled with digoxigenin-dUTP or biotin-dUTP using a nick translation kit (Roche), denatured (95°C, 5 min) and pre-annealed (37°C, 45 min) with Cot-1 DNA and salmon sperm DNA in hydridisation buffer (50% formamide, 20% dextran sulphate, 2X SSC, 1 mg/ml BSA) immediately before hybridization. Digoxigenin FISH signals were amplified using sheep anti-digoxigenin fluorescein fab fragments (1:50, Roche 11207741910) and fluorescein rabbit anti-sheep antibodies (1:100, Vector Labs FI-6000); biotin probes were detected using streptavidin-555 (1:1000, Molecular Probes). For single FISH experiments, digoxigenin labelling was used; for double DNA-FISH pairs of probes with different labels were mixed immediately prior to addition to the coverslip for hybridisation. DNA was counterstained with 4⍰,6-diamidino-2-phenylindole (DAPI). Coverslips were washed and mounted in Prolong Gold (Thermo Fisher Scientific). Confocal images of neuronal nuclei were acquired using a Leica SPE3 confocal microscope for lamina association, or an SP8 confocal microscope for double DNA-FISH. Images were analyzed using Fiji software. Probe coordinates were identified using the 3D Objects Counter tool on hyperstacks of individual nuclei (ensuring only 1 or 2 foci per cell). For double DNA-FISH analysis, the separation of the probe coordinates (distance AB) from each channel were calculated using the formula:

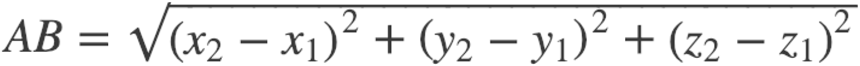

For measurements of probe to nuclear periphery, the edge of the nucleus was identified using the DAPI staining.

### Chromatin Immunoprecipitation

Chromatin immunoprecipitation (ChIP) experiments were performed as described previously (Policarpi et al., 2017) with some modifications. To crosslink proteins with DNA, the medium was removed from neuronal cultures, and crosslinking buffer (0.1 M NaCl, 1 mM EDTA, 0.5 mM EGTA and 25 mM HEPES-KOH, pH 8.0) containing 1% formaldehyde was added for 10 min at room temperature. The cross-linking reaction was stopped by adding glycine to a final concentration of 125 mM. Cells were rinsed three times with ice-cold PBS containing protease inhibitor cocktail and 1 mM PMSF, collected by scraping and centrifuged at 3000 rpm at 4°C for 10 minutes. Cell pellets were transferred to 1.5 ml tubes and lysed with 20 cell pellet volumes (CPVs) of buffer 1 (50 mM HEPES-KOH, pH 7.5, 140 mM NaCl, 1 mM EDTA, pH 8.0, 10% glycerol, 0.5% NP-40, 0.25% Triton X-100 and complete protease inhibitor cocktail) for 10 min at 4 °C. Nuclei were pelleted by centrifugation at 3000 rpm for 10 min at 4 °C, incubated with 20 CPVs of buffer 2 (200 mM NaCl, 1 mM EDTA, pH 8.0, 0.5 mM EGTA, pH 8.0, 10 mM Tris-HCl, pH 8.0, and complete protease inhibitor cocktail) for 10 min at RT and re-pelleted. 4 CPVs of buffer 3 (1 mM EDTA, pH 8.0, 0.5 mM EGTA, pH 8.0, 10 mM Tris-HCl, pH 8.0, and complete protease inhibitor cocktail) were added to the nuclei, and sonication was carried out by applying 20 pulses, 30 seconds each, at 30 seconds intervals. Insoluble materials were removed by centrifugation at 14000 rpm for 10 min at 4 °C, the supernatant was transferred to a new tube and the final volume of the nuclear lysate was adjusted to 1 ml by adding buffer 3 supplemented to give 150 m M NaCl, 1% Triton-X, 0.1% sodium deoxycholate in the final chromatin sample. 50 μl of the 1 ml chromatin samples was saved for an Input, while the remaining fraction was incubated with 5 μg Rad21 (ab992) antibody and 50 μl Dynabeads (Thermo Fisher Scientific; washed once) and rotated overnight at 4°C. Beads were pelleted and washed with: low-salt buffer (0.1% SDS, 1% Triton X-100, 2 mM EDTA, 20 mM Tris-HCl, pH 8.0, 150 mM NaCl), high-salt buffer (0.1% SDS, 1% Triton X-100, 2 mM EDTA, 20 mM Tris-HCl, pH 8.0, 500 mM NaCl) and LiCl buffer (0.25 M LiCl, 1% IGEPAL CA630, 1% deoxycholic acid (sodium salt), 1 mM EDTA, 10 mM Tris, pH 8.1) and twice with TE buffer (10 mM Tris-HCl, pH 8.0, 1 mM EDTA). For each wash, the beads were incubated for 10 min at 4 °C while rotating. The immunoprecipitated DNA was eluted by adding elution buffer (0.1 M NaHCO_3_ pH 8.0, 1% SDS) and incubating at 65°C, 5 min and then rotating at RT for 15 min. Crosslinking was reversed by adding 10 μl 5M NaCl and incubating the samples at 65°C overnight. DNA was purified using PCR purification columns (Qiagen), quantified using the Qubit high sensitivity assay and subjected to qPCR using the same amount of DNA in immunoprecipitated and input PCRs. Primer sequences are shown in **Supplementary Table S3**. The protocadherin HS5 region was used as a positive control (Monahan et al., 2012).

### 4C-seq

4C-seq experiments were performed as described previously (Sofueva et al., 2013). To crosslink proteins with DNA, the medium was removed from neuronal cultures, and crosslinking buffer (0.1 M NaCl, 1 mM EDTA, 0.5 mM EGTA and 25 mM HEPES-KOH, pH 8.0) containing 1% formaldehyde was added for 10 min at room temperature. The cross-linking reaction was stopped by adding glycine to a final concentration of 125 mM. Cells were rinsed three times with ice-cold PBS containing protease inhibitor cocktail and 1 mM PMSF, collected by scraping and centrifuged at 3000 rpm at 4°C for 10 minutes. Cell pellets were lysed in 10 ml lysis buffer (10 mM Tris pH 8.0, 10 mM NaCl, 0.2% NP40 supplemented with protease inhibitor cocktail and 1 mM PMSF) on ice for 20 min. Nuclei were collected by centrifugation (1800 rpm, 5 min, 4°C), resuspended in 1.2X DpnII buffer and transferred to Protein LoBind tubes. SDS was added to 0.3% final concentration and nuclei were incubated 1h at 37°C in thermomixer shaking at 900 rpm (30s on, 30s off). Triton-20 was added to 2% final concentration and nuclei were incubated 1h 37°C in a thermomixer shaking at 900 rpm (30s on, 30s off). 750 Units of DpnII (NEB) was added and incubated overnight at 37°C in a thermomixer shaking at 900 rpm (30s on, 30s off). The next day, the DpnII buffer was replaced with fresh 1.2X DpnII buffer supplemented with 0.3% SDS and 2% Triton and another 750 Units of DpnII and incubated overnight at 37°C in thermomixer shaking at 900 rpm (30s on, 30s off). Samples of undigested and DpnII-digested DNA was reverse crosslinked and run on a gel to confirm that most DNA fragments were <3 kb after digestion. Nuclei were centrifuged (1800 rpm, 3 min) and washed twice with 1X T4 DNA ligase buffer before resuspending in 100 μl 1X T4 DNA ligase buffer with 1600 Units T4 DNA ligase (NEB). In nucleo ligation was carried out overnight at 16°C without shaking before confirming that high molecular weight products were obtained. Samples were then reverse crosslinked in the presence of proteinase K overnight at 65°C before phenol:chloroform extraction and ethanol precipitation. DNA was quantified using Qubit high sensitivity assays (Thermo Fisher Scientific)6-10 μg of DNA was digested with 120 Units Csp6I enzyme (Thermo Fisher Scientific) [3-5 Csp6I digests per sample] overnight at 37°C in thermomixer shaking at 900 rpm (30s on, 30s off). After confirmation that Csp6I-digested products are <3kb, Csp6I was heat inactivated at 65°C for 20 min before phenol:chloroform extraction and ethanol precipitation. DNA was resuspended in 6 ml total volume to allow proximity ligation by 1600 Units T4 DNA ligase overnight at 16°C. Samples were purified by phenol:chloroform extraction and ethanol precipitation, followed by PCR purification columns (Qiagen), before quantitation using with Qubit high sensitivity assays (Thermo Fisher Scientific). 4C-seq libraries were generated using Expand Long Template polymerase (Roche) and primers designed using the 4C-seq primer database (van de Werken et al., 2012) (**Supplementary Table S2**). Forward primers were generated with the Illumina p1 sequence (AATGATACGGCGACCACCGAGATCTACACTCTTTCCCTACACGACGCTCTTCCGATCT), a two-nucleotide barcode to allow multiplexing of samples, and then the primer sequence. Reverse primers were generated with the Illumina p2 CAAGCAGAAGACGGCATACGAGATCGGTCTCGGCATTCCTGCTGAACCGCTCTTCCGATCT). 6-10 PCRs were set up per sample to generate library diversity. PCRs were run using the following program: 3 min 94°C; then 29 cycles of 10s 94°C, 1 min 55°C, 3 min 68°C; then 10 min 68°C. PCR products were purified using the High Pure PCR product purification kit (Roche). Libraries were quantified with Qubit high sensitivity assays, assessed using the Agilent Tapestation, and run on an Illumina MiSeq (MiSeq Reagent Kit v3, 150-cycle). 4C-seq data analysis and normalization was performed as described (van de Werken et al., 2012).

### CRISPR-Cas9 vectors

Single guide RNAs were designed towards the putative *Bdnf* enhancer using http://crispr.mit.edu/. The sequences of the guide RNAs that we used throughout this study are (last 3 nucleotides are PAM):

Enh^g1^ GGATTGTTTGGACTTACTCT

Enh^g2^ GTTTTGTCAAGTGTGGGAGC

The backbones for the BPK1520 vector used to express the guide RNA (U6-BsmBIcassette-Sp-sgRNA) was a gift from Keith Joung (Addgene 65777). Annealed oligos composing the different guide RNAs were cloned into the BsmBI site of U6-BsmBIcassette-Sp-sgRNA. The CRISPRi repressor dCas9-KRAB-MECP2 (Yeo et al., 2018) was a gift from Alejandro Chavez and George Church (Addgene 110821). The backbones for the lentiviral vector pLV hU6-sgRNA hUbC-dCas9-KRAB-T2a-GFP and pLV hU6-sgRNA hUbC-dCas9-KRAB-T2a-puro (Thakore et al., 2015) were a gift from Charles Gersbach (Addgene 71236, 71237) and the same guides targeting the *Bdnf* enhancer were cloned into the BsmBI site.

Bdnf or control EV overexpression vectors were a gift from Christian Rosenmund (Sampathkumar et al., 2016).

### Lentiviral production

10 μg of the transfer vector pLV hU6-sgRNA hUbC-dCas9-KRAB-T2a-puro (Empty, or containing Enh^g1^ or Enh^g2^) was transfected into each 10 cm dish of HEK293T cells together with the packaging vectors psPax2 (7.5 μg) and pCMV-VSV-G (5 μg) using PEImax (67.5 μg; Polysciences) or Lipofectamine-2000 (50 μl; Thermo Fisher Scientific) in Opti-MEM (Thermo Fisher Scientific). The media was changed after 4h to HEK293T media (DMEM plus 10% fetal bovine serum, 2 mM L-glutamine, 1X penicillin/streptomycin) supplemented with 1% BSA to improve viral stability. The media containing viral supernatant was harvested 48h and 72h later. Viral supernatant from all plates was combined, passed through 0.45 μm syringe filters and concentrated using PEG precipitation or ultracentrifugation. For PEG precipitation, PEG was mixed with the media to 10% final concentration and incubated overnight at 4°C. Samples were centrifuged 2500 rpm, 20 min and the supernatant discarded. For ultracentrifugation, media containing viral particles was ultracentrifuged at 24000 rpm, 2h, 4°C in a Beckman Optima XPN-80 Ultracentrifuge. The pellets were resuspended in Neurobasal media (Thermo Fisher Scientific) at a 200X concentration.

### Immunofluorescence and clustering analysis

Cells grown on coverslips were washed in PBS and then fixed in 4% PFA (TAAB, 20 min, RT). Cells were washed in PBS (3 times 3 min, RT), permeabilised in 0.3% Triton-X in PBS (10 min, RT) and then blocked in 5% goat serum, 5% fetal bovine serum in 1X PBS (1h, RT). Primary antibody incubations took place in a humid chamber at 4°C overnight with the following antibodies: chicken anti-GFP (Abcam ab13970 1:2000), mouse anti-mCherry (ab125096 1:1000). Cells were washed in PBS (3 times 3 min, RT) before amplification and detection using goat anti-chicken AlexaFluor-488 and donkey anti-mouse AlexaFluor-555 (1:1000, Molecular Probes). Coverslips were washed and mounted in Prolong Gold (Thermo Fisher Scientific). DNA was counterstained with 4⍰,6-diamidino-2-phenylindole (DAPI). Coverslips were blinded and confocal images of neuronal nuclei were acquired using a Leica SPE3 confocal microscope.

Clustering of neuronal cells was analysed in Fiji using maximal z projections of the DAPI channel (each image was of a single neuronal cluster and its surrounding cells; if the edge of another cluster was in the image this was removed before processing). After applying a Gaussian blur filter (Sigma 4.0) to even out the signal, we used the ‘Find Maxima’ tool to identify each nucleus. The XY coordinates were inputted into R and used to compute the distance between every point and every other point, before the median per image was calculated and samples were deblinded.

### Dendritogenesis assays

Assays were carried out as described previously (Crepaldi et al., 2013). Briefly, 2-3 hours after plating in 24-well plates, mouse cortical neurons were transfected using Optimem containing 375 ng dCas9-KRAB-MECP2 DNA and 125 ng BPK1520 (Empty, or containing guides targeting the *Bdnf* enhancer) and a GFP expression vector (200 ng pBIRD (Supplementary Fig. 2) or 500 ng pCIG vector (EV or p*Bdnf*); Figure 3) and 0.8-1.5 μl Lipofectamine 2000 (Life Technologies). After 2 hours, the medium was replaced with culture media containing 0.33X B27 (serum starve conditions) with or without 50 mM KCl. Cells were cultured for 48 hours followed by immunostaining with anti-GFP (Abcam ab13970, 1:2000). Coverslips were blinded before images of GFP-transfected non-overlapping neurons were obtained using a Zeiss Axio Imager microscope and analyzed in Fiji. For quantification of total dendritic length and Sholl analysis we used the Simple Neurite tracer plugin, and then samples were deblinded.

### Data and code availability statement

4C-seq data is available through GEO (accession number GSE…). Codes were all previously published.

