## Supplementary Figures and Tables for "A novel enhancer that regulates *Bdnf* expression in developing neurons"

### Figure S1

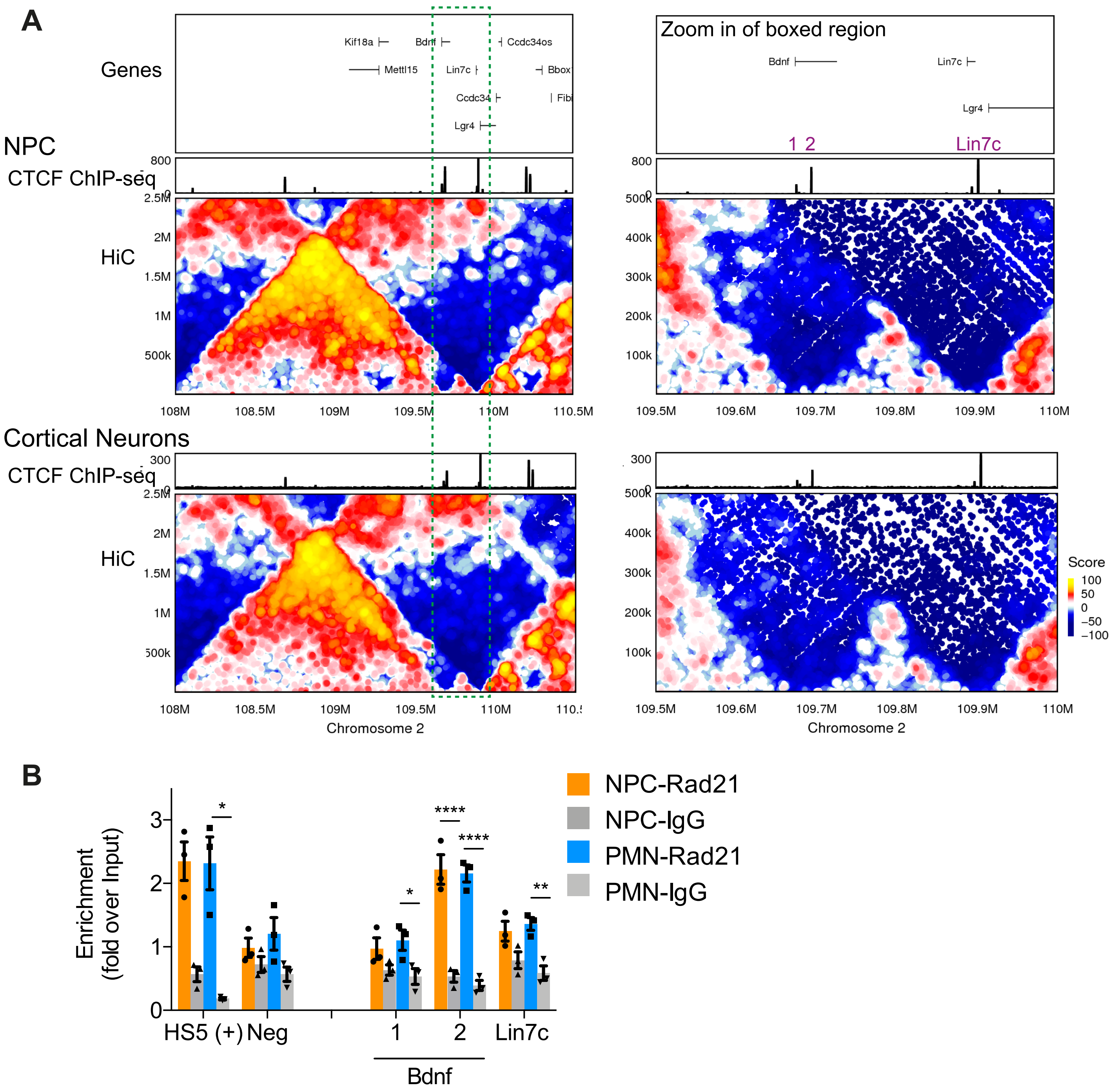
A
Genes
Kif18a
Bdnf
Ccdc34os
Mettl15
Lin7c
Bbox
Ccdc34
Lgr4
NPC
CTCF ChIP-seq
HiC
108M
108.5M
109M
109.5M
110M
110.5M
Zoom in of boxed region
1 2
Lin7c
109.5M
109.6M
109.7M
109.8M
109.9M
Cortical Neurons
Chromosome 2
Score
100
50
0
−50
−100
B
Enrichment (fold over Input)
HS5 (+)
Neg
1
2
Lin7c
Bdnf
NPC-Rad21
NPC-IgG
PMN-Rad21
PMN-IgG

### Figure S2

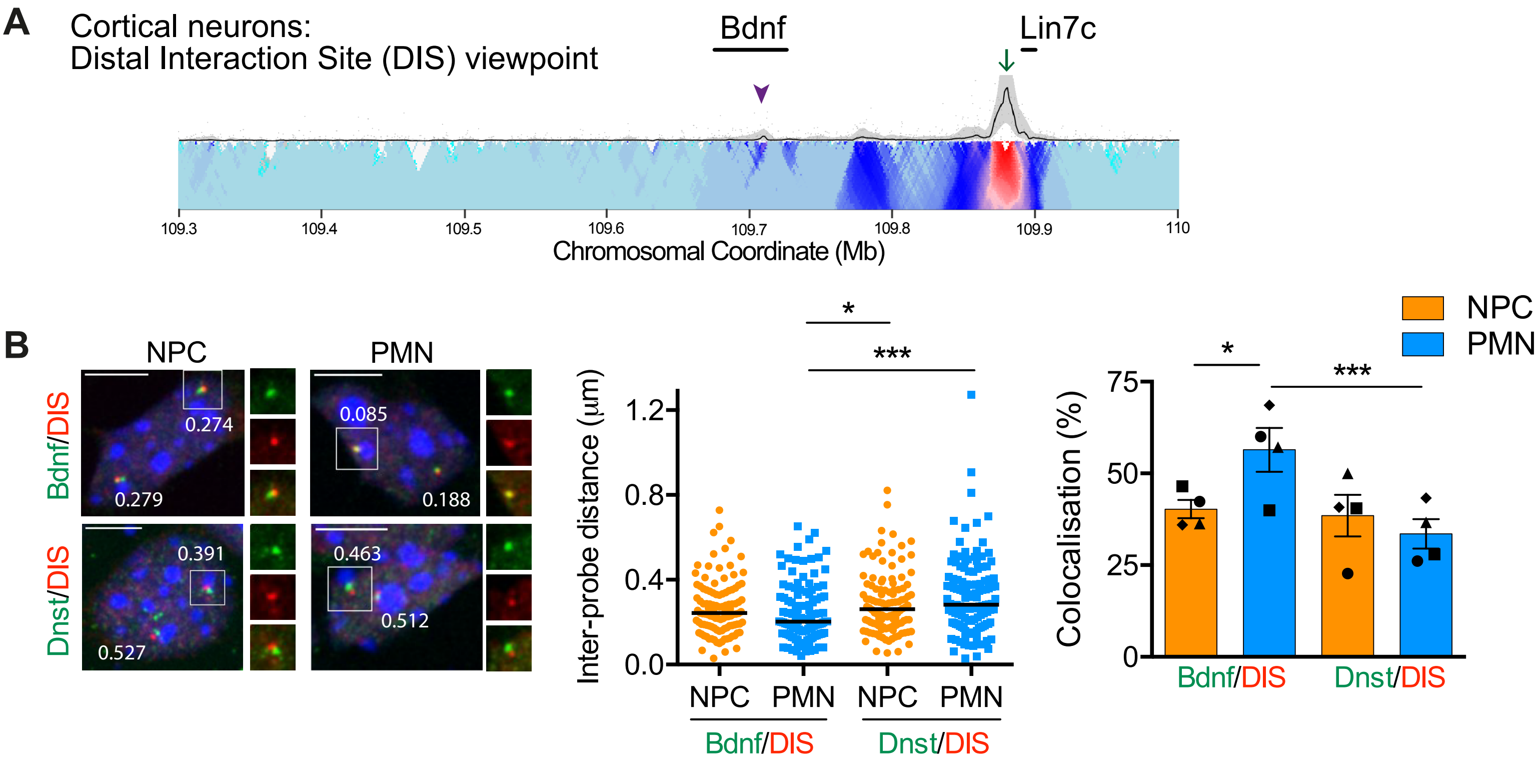
A
Cortical neurons:
Distal Interaction Site (DIS) viewpoint
Bdnf
Lin7c
109.3
109.4
109.5
109.6
109.7
109.8
109.9
110
Chromosomal Coordinate (Mb)
B
NPC
PMN
Bdnf/DIS
Dnst/DIS
0.274
0.279
0.085
0.188
0.391
0.527
0.463
0.512
Inter-probe distance (μm)
1.2
0.8
0.4
0.0
NPC
PMN
NPC
PMN
Bdnf/DIS
Dnst/DIS
*
***
NPC
PMN
Colocalisation (%)
75
50
25
0
*
***
Bdnf/DIS
Dnst/DIS

#### Figure S3

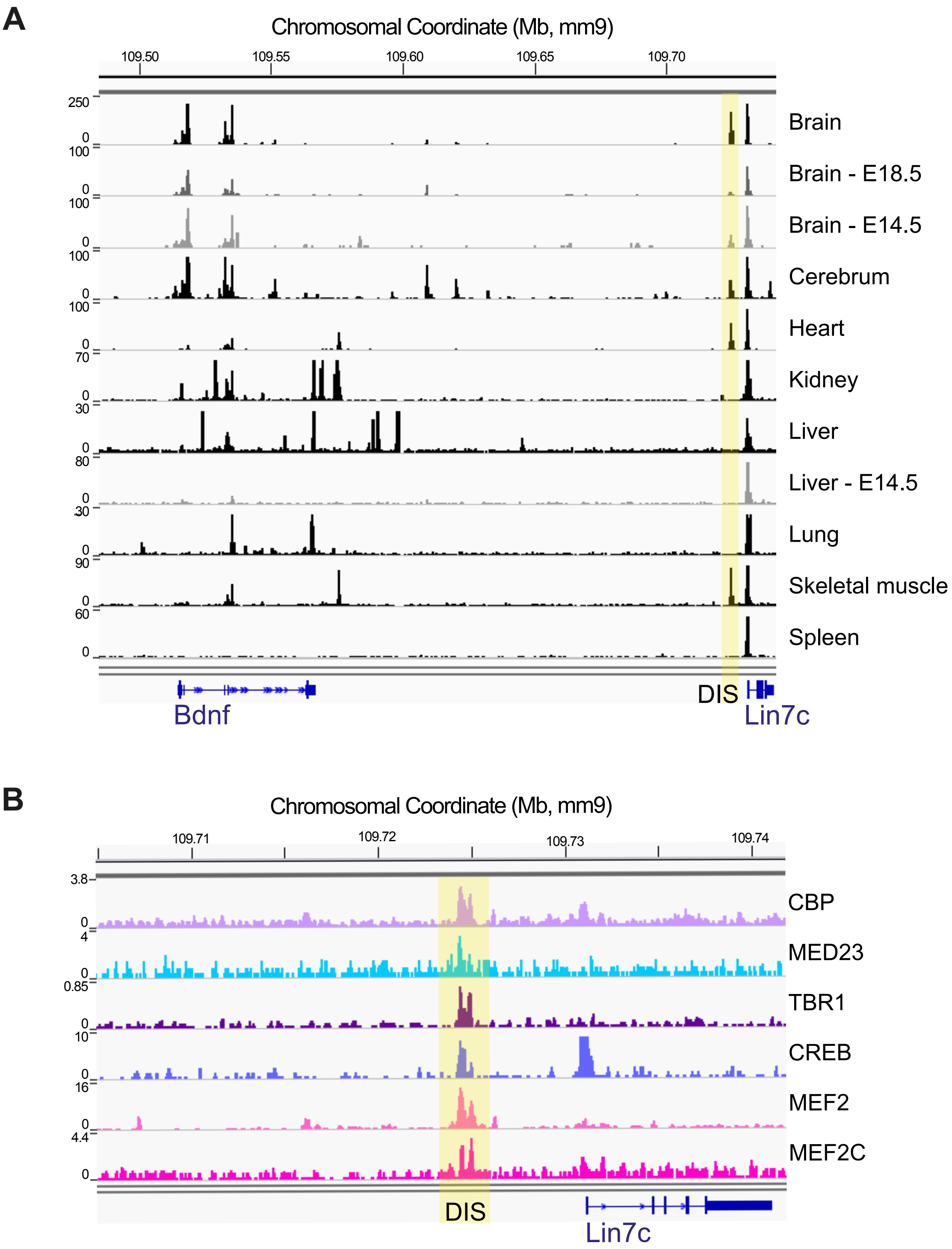
A
Chromosomal Coordinate (Mb, mm9)
109.50
109.55
109.60
109.65
109.70
250
0
100
0
100
0
100
0
100
0
70
0
30
0
80
0
30
0
90
0
60
0
Brain
Brain - E18.5
Brain - E14.5
Cerebrum
Heart
Kidney
Liver
Liver - E14.5
Lung
Skeletal muscle
Spleen
DIS
Bdnf
Lin7c
B
Chromosomal Coordinate (Mb, mm9)
109.71
109.72
109.73
109.74
3.8
0
4
0
0.85
0
10
0
16
0
4.4
0
CBP
MED23
TBR1
CREB
MEF2
MEF2C
DIS
Lin7c

#### Figure S4

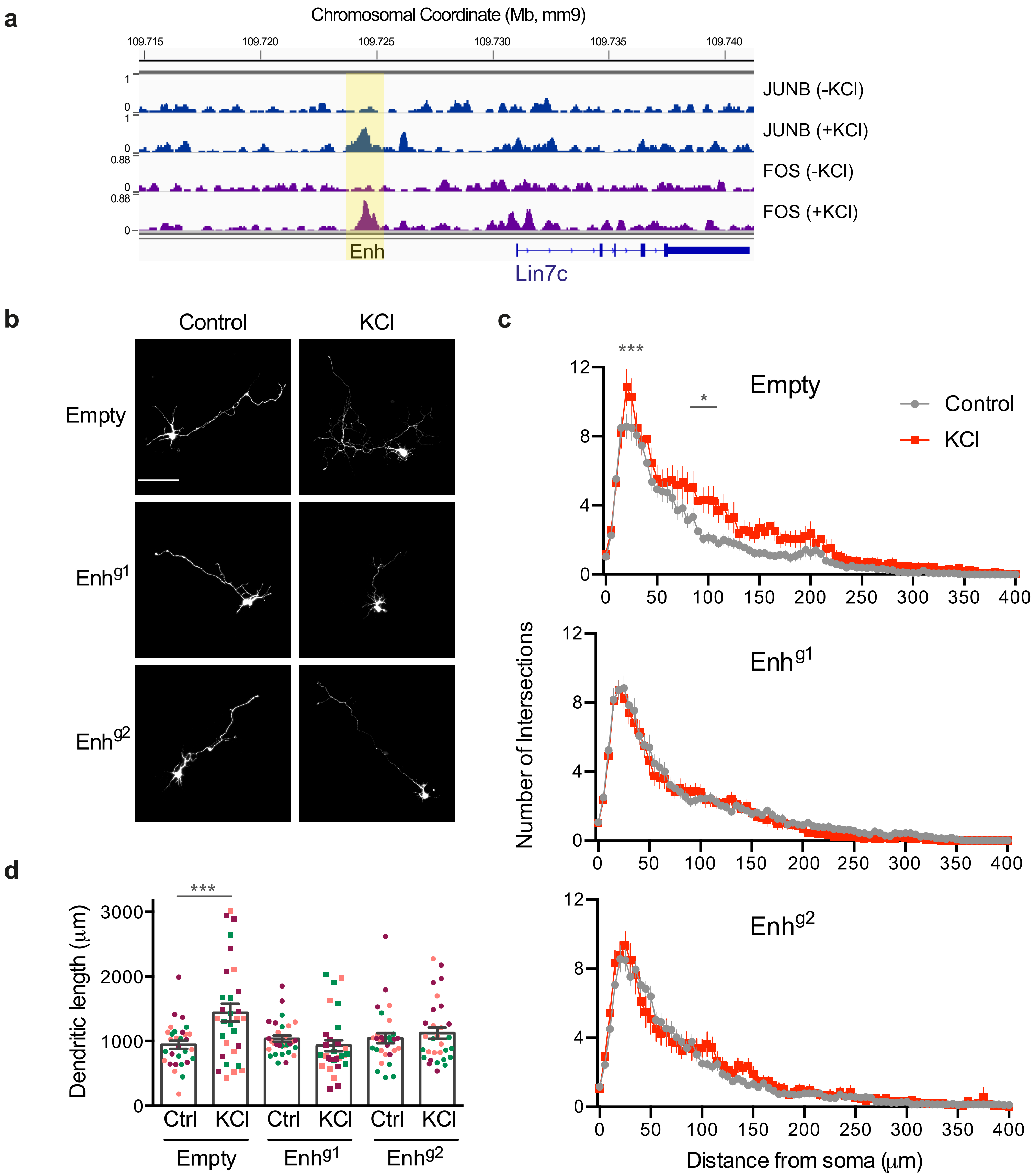
a
Chromosomal Coordinate (Mb, mm9)
109.715
109.720
109.725
109.730
109.735
109.740
JUNB (-KCl)
JUNB (+KCl)
FOS (-KCl)
FOS (+KCl)
Enh
Lin7c
b
Control
KCl
Empty
Enhg1
Enhg2
c
Empty
Control
KCl
Enhg1
Enhg2
Number of Intersections
Distance from soma (μm)
d
Dendritic length (μm)
Ctrl
KCl
Empty
Enhg1
Enhg2

#### Supplementary Figure Legends

**Supplementary Figure S1. *Bdnf* resides in a small sub-TAD containing the gene and a downstream intergenic region.**

**A)** HiC and CTCF ChIP-seq data from NPCs and cortical neurons around the *Bdnf* locus discriminates a sub-TAD containing *Bdnf*, a downstream intergenic region and the *Lin7c* gene. This sub-TAD is at the 3' end of a larger TAD, and its boundaries are CTCF-positive. Right panel; zoom of left panel. Purple numbers indicate sites positive for CTCF in the ChIP-seq data, which were used for Rad21 ChIP-qPCR analysis.

**B)** ChIP-qPCR of the cohesin subunit Rad21 in NPC and PMN confirms Rad21 binding at the boundaries of the *Bdnf* sub-TAD. Immunoglobulin G (IgG) was used as a negative control. The protocadherin HS5 region was used as a positive control (Monahan et al., 2012). Bars with error bars represent mean ± SEM, and points show results from different biological replicates ($n$ = 3). $^{*}p < 0.05$, $^{**}p < 0.01$, $^{***}p < 0.001$, $^{****}p < 0.0001$, two-way ANOVA with Sidak's multiple comparisons test. HS5: PMN-Rad21 vs. PMN-IgG $p$=0.0455. Bdnf-CTCF1: PMN-Rad21 vs. PMN-IgG $p$=0.0384. Bdnf-CTCF2: NPC-Rad21 vs. NPC-IgG $p$<0.0001; PMN-Rad21 vs. PMN-IgG $p$<0.0001. Lin7c-CTCF: PMN-Rad21 vs. PMN-IgG $p$=0.0052.

**Supplementary Figure S2. *Bdnf* forms a chromatin loop with an intergenic region and the Lin7c gene.**

**A)** 4C-seq contact profiles in cortical neurons from a viewpoint designed upstream of Lin7c, at the distal interaction site. Exon I or putative enhancer viewpoints. Image shows a single representative 4C-seq experiment (from $n$=2; profiles from biological replicates were very similar) represented by the median normalized 4C-seq coverage in a sliding window of 5 kb (top) and a multi-scale domainogram indicating normalized mean coverage in windows ranging between 2 and 50 Kb. Green arrow indicates viewpoint. Arrowhead indicates interaction with the *Bdnf* gene.

**B)** Double DNA-FISH of the DIS (Distal Interacting Site) with either the *Bdnf* gene or an equidistant region downstream with reciprocal labelling to that shown in **Fig. 2b**. Left panel, representative images of confocal sections of double DNA FISH in NPCs and PMNs. Nuclei were stained with DAPI (blue). Middle panel, scatter dot plot of interprobe distance measurements (NPC, orange; PMN, blue). Solid black lines denote means. $^{****}p < 0.001$, One-way ANOVA with Dunn's multiple comparisons (two-tailed). $n$ = 111 (Bdnf/DIS-NPC), 117 (Bdnf/DIS-PMN), 114 (Dnst/DIS-NPC), 126 (Dnst/DIS-PMN) foci across 4 biological replicates. Probe labelling denoted in coloured font. Bdnf/DIS-PMN vs. Dnst/DIS-NPC $p$=0.0403; Bdnf/DIS-PMN vs. Dnst/DIS-PMN $p$=0.0005. Right panel, colocalisation (defined as an inter-probe distance of 225nm or less) of FISH signals in double DNA FISH experiments performed in NPCs and PMNs. $^{***}p<0.001$, $^{*}p<0.05$, Fisher's exact test (two-tailed). Bdnf/DIS-NPC vs. Bdnf/DIS-PMN $p$=0.0123; Bdnf/DIS-PMN vs. Down/DIS-PMN $p$=0.0002.

**Supplementary Figure S3. *Bdnf*-interacting intergenic region displays hallmarks of an enhancer.**

**A)** DNaseI hypersensitivity across the *Bdnf* locus in the indicated tissues and developmental stages (black, adult; grey, embryonic) shows chromatin accessibility at the distal interacting site (DIS) and the *Bdnf* gene in embryonic and adult brain and cerebrum but not other tissues.
**B)** CBP, Mediator and transcription factors bind to the DIS. Published ChIP-seq datasets (see **Supplementary Table S1**) mapped to mm9.

**Supplementary Figure S4. CRISPR inhibition of the putative Bdnf enhancer prevents activity-dendritogenesis.**

**A)** FOS and JUNB ChIP-seq in cortical neurons minus and plus KCl (2h) show activity-dependent recruitment to the *Bdnf* enhancer. ChIP-seq data (Malik et al., 2014) mapped to mm9 genome.
**B)** Representative images of cortical neurons transfected with a GFP (pBIRD) in combination with dCas9-KRAB-MECP2 and an expression vector for guide RNAs (Empty or targeting the putative enhancer region ($Enh^{g1}$, $Enh^{g2}$)). Cells were maintained under basal or depolarizing (KCl, 50 mM) conditions for 48 hr, followed by GFP immunostaining. Scale bar, 100 µm.
**C)** Sholl analysis of the dendritic processes of 30 neurons per treatment (10 per biological replicate). For each distance point, the mean number of intersections ± SEM is shown. $^{*}p < 0.05$, $^{**}p < 0.01$, $^{***}p < 0.001$, $^{****}p < 0.0001$, two-way ANOVA with Sidak's multiple comparisons test. Empty: Control vs. KCl 20 µm from soma $p$=0.0009, mean diff -2.267, 95% CI of diff -4.056 to -0.4775; 80 µm from soma $p$=0.0280, mean diff -1.867, 95% CI of diff -3.656

to -0.07752; 95 µm from soma $p$=0.0013, mean diff -2.233, 95% CI of diff -4.022 to -0.4442; 100 µm from soma $p$=0.0017, mean diff -2.200, 95% CI of diff -3.989 to -0.4109; 105 µm from soma $p$=0.0023, mean diff -2.167, 95% CI of diff -3.956 to -0.3775; 110 µm from soma $p$=0.0216, mean diff -1.900, 95% CI of diff -3.689 to -0.1109; 115 µm from soma $p$=0.0216, mean diff -1.900, 95% CI of diff -3.689 to -0.1109.

**D)** Quantification of the total length of the dendritic processes of neurons analysed in (b). Bars show mean ± SEM; points show each data for each neuron coloured by biological replicate ($n$=3). $^{***}p < 0.001$, two-way ANOVA with Sidak's multiple comparisons test. Empty: Ctrl vs. KCl $p$=0.0003, mean diff -497.1, 95% CI of diff -797.0 to -197.2.

**Supplementary Tables**

**Supplementary Table S1. Published datasets used in this study**

| Data | Cell type | Reference | Accession number |
|---|---|---|---|
| **HiC** | NPC | (Bonev et al., 2017) | GSE96107 |
| | CN | (Bonev et al., 2017) | GSE96107 |
| **CTCF ChIP-seq** | NPC | (Bonev et al., 2017) | GSE96107 |
| | CN | (Bonev et al., 2017) | GSE96107 |
| **H3K4me1 ChIP-seq** | CN – KCl | (Policarpi et al., 2017) | GSE75191 |
| | CN + KCl | (Policarpi et al., 2017) | GSE75191 |
| **H3K27ac ChIP-seq** | CN – KCl | (Policarpi et al., 2017) | GSE75191 |
| | CN + KCl | (Policarpi et al., 2017) | GSE75191 |
| **CBP ChIP-seq** | CN - Reelin | (Telese et al., 2015) | GSE66710 |
| **CREB ChIP-seq** | CN - Reelin | (Telese et al., 2015) | GSE66710 |
| **MED23 ChIP-seq** | CN - Reelin | (Telese et al., 2015) | GSE66710 |
| **MEF2C ChIP-seq** | CN - Reelin | (Telese et al., 2015) | GSE66710 |
| **MEF2 ChIP-seq** | CN - Reelin | (Telese et al., 2015) | GSE66710 |
| **TBR1 ChIP-seq** | E15.5 cortex | (Notwell et al., 2016) | GSE71384 |
| **JUNB ChIP-seq** | CN – KCl | (Malik et al., 2014) | GSE60192 |
| | CN + KCl | (Malik et al., 2014) | GSE60192 |

| | | | |
|---|---|---|---|
| **FOS ChIP-seq** | CN – KCl | (Malik et al., 2014) | GSE60192 |
| | CN + KCl | (Malik et al., 2014) | GSE60192 |
| **DNase HS** | Tissues | ENCODE | |
| **GRO-seq** | CN - Reelin | (Telese et al., 2015) | GSE66710 |
| | CN + Reelin | (Telese et al., 2015) | GSE66710 |

**Supplementary Table S2. 4C-seq primers.**

| Bait | Primer sequence |
|---|---|
| **Exon1_F** | CCGGACATCTGCCTAGGATC |
| **Exon1_R** | CCCTCCTATCCTAAGAATGC |
| **Enhancer_F** | TGCTACATGTGGTAAAGATC |
| **Enhancer_R** | GCTTAGCACCTATGCTCAGT |

Primers sequences were taken from the mouse database (van de Werken et al., 2012). The primers were synthesized with sequences for Illumina sequencing:

F: AATGATACGGCGACCACCGAGATCTACACTCTTTCCCTACACGACGCTCTTCCGATCT

R: CAAGCAGAAGACGGCATACGAGATCGGTCTCGGCATTCCTGCTGAACCGCTCTTCCGATCT)

with a barcode (GG, AC, AG, CC) upstream of the forward primer sequence.

**Supplementary Table S3. Primers used for real time PCR analysis of reverse transcription and chromatin immunoprecipitation products**

| Name | Sequence (5’-3’) | Application |
|---|---|---|
| **Bdnf_allRNA_F1** | GGCCCAACGAAGAAAACCAT | qRT-PCR |
| **Bdnf_allRNA_R1** | GTTTGCGGCATCCAGGTAAT | qRT-PCR |
| **Bdnf_putENH-A_F** | GGCTGTACACTTCCTCTCCA | qRT-PCR |
| **Bdnf_putENH-A_R** | GCTTTCCCCATTCCTTGACT | qRT-PCR |
| **Bdnf_putENH-B_F** | AGCTGTGCCTTATATGTACTTCA | qRT-PCR |
| **Bdnf_putENH-B_R** | ACCCTGCTTCTCTTGTTCTCT | qRT-PCR |
| **Bdnf_Universal_R** | GCCTTCATGCAACCGAAGTA | qRT-PCR |
| **Bdnf_Ex1_F** | TGCATCTGTTGGGGAGACAA | qRT-PCR |

| | | |
|---|---|---|
| **Bdnf_Ex2_F** | CCATTCAGCACCTTGGACAG | qRT-PCR |
| **Bdnf_Ex3_F** | GGATGCTTCATTGAGCCCAG | qRT-PCR |
| **Bdnf_Ex4_F** | AGCTGCCTTGATGTTTACTTTGA | qRT-PCR |
| **Bdnf_Ex5_F** | TTTCTAGCTTTGTGGTGCGG | qRT-PCR |
| **Bdnf_Ex6_F** | ATCCGAGAGCTTTGTGTGGA | qRT-PCR |
| **Bdnf_Ex7_F** | CTGAAAGGGTCTGCGGAACT | qRT-PCR |
| **Bdnf_ex8_F** | ATCCCAGGAGAAAGGCTGTG | qRT-PCR |
| **Bdnf_Ex9_F** | TTACAAGCAGATGGGCCACA | qRT-PCR |
| **Lin7c_Ex3-4_F** | CCCTGAAGTGAGAGCCAATG | qRT-PCR |
| **Lin7c_Ex3-4_R** | CTGTCAGCAATTCCACCTGG | qRT-PCR |
| **NeuN_F** | CCAGGCACTGAGGCCAGCACACAGC | qRT-PCR |
| **NeuN_R** | CTCCGTGGGGTCGGAAGGGTGG | qRT-PCR |
| **bactin_F** | TCTTTGCAGCTCCTTCGTTG | qRT-PCR (HK) |
| **bactin_R** | ACGATGGAGGGGAATACAGC | qRT-PCR (HK) |
| **bactin_ex-int_F** | GATATCGCTGCGCTGGTC | qRT-PCR |
| **bactin_ex-int_R** | CATCGATCCCCAAGAAAACC | qRT-PCR |
| **Bdnf_CTCF1_F** | TTTGGTCCCCTCATTGAGCT | ChIP |
| **Bdnf_CTCF1_R** | TTCTTTGCGGCTTACACCAC | ChIP |
| **Bdnf_CTCF2_F** | GCTAGGAAGGTAGAGGGTGC | ChIP |
| **Bdnf_CTCF2_R** | AGCCAAAATTCCGAACCGAG | ChIP |
| **Lin7c_CTCF1_F** | CATTTGCTGCCAGTGAAGGA | ChIP |
| **Lin7c_CTCF1_R** | CTGTCAGCAATTCCACCTGG | ChIP |
| **HS5_F** | GCCATGGAGATTTTCTTTACATGA | ChIP |
| **HS5_R** | TGGCAGATGGAACCACTTTTTA | ChIP |
| **Neg_F** | GGACAATTCAACCGAGGAAA | ChIP |
| **Neg_R** | TGAACTGGTTTGGTGTGCTC | ChIP |
